# Shapes of synaptic protein distributions distinguish brain architectures

**DOI:** 10.64898/2025.12.19.695447

**Authors:** Martin Rehn, Mélissa Cizeron, Zhen Qiu, Seth G. N. Grant, Erik Fransén

## Abstract

Neuronal activity is the result of the orchestrated actions of multitudes of synapses, acting and evolving not as individuals, but as populations. We therefore collected, for millions of synapses, a functionally relevant proxy for their strengths. Namely, we measured postsynaptic protein content by deploying a fluorescent PSD95 variant. Then we summarized, by statistical moments, the shapes of the distributions of protein contents, over local synaptic populations. In this way we explored the hypothesis that such collective properties inform on brain architectures.

Our measurements cover complete parasagittal sections of the mouse brain, in animals one week to 18 months old. We identified a hierarchical organization of regions along the anterior–posterior axis, with three main clusters of divergent synaptic population shapes. One includes telencephalic regions, one is centered in the midbrain and hindbrain, and one comprises mainly the thalamus and the cerebellum. The structure emerging from our approach aligns with discoveries in studies of regional patterns of cellular gene expressions. Our results suggest that synaptic populations are dynamically regulated over the lifespan. Regions which at three months of age have thinner tails largely conserve their distribution shapes later in life, whereas heavy tails in other regions strikingly grow ever more so.

## Introduction

Animal behavior is due to activity in neuronal populations, controlled by synaptic transmission. Neurons in the central nervous system typically receive at least thousands of synapses, which has two consequences. Firstly, the bulk of information processing almost certainly occurs on the synaptic, rather than on the neuronal level, as further evidenced by the complexity of the synaptic protein machinery, which is most prominent in the postsynaptic density. Secondly, neuronal activity being mediated in general by the integration of many inputs hints that it is the collective actions of populations of synapses that are the building blocks of brain function. Synaptic populations change dramatically during early development, more subtly in the adult, and have been generally thought to also change subtly in the aging animal. Synaptic processes: the creation, modification, and removal of synapses underpin learning and behavioral adaptation, as well as regulate homeostasis, notably the preservation of excitation–inhibition balance. The dynamics and effects of these forces have been observed for instance in monocular enucleation (Barnes et al., 2017) and in enriched environments (Wegner et al., 2022). Drifts in synapses over the sleep and wakefulness cycle (de Vivo et al., 2017) have furthermore been suggested to be consistent with a specific regulatory model of opposing forces: learning stretches the distribution, making the synaptic populations more diverse, while homeostatic plasticity shifts the distribution back towards a set point (Sammons et al., 2018). As an example of such potential balancing, experiments affecting spines on dentate granule neurons impacted presynaptic activity, plasticity induction, and TNF and TNF-associated receptors (Rößler et al., 2023, 2024), while a skewed distribution shape was preserved. These observations illuminate the balance between small and large synapses, how it changes over the lifespan (Parajuli et al., 2020), and imply that the distributions across populations carry important information.

Studies of species populations in ecology, as well as in genes, mRNA, and of proteins in cell biology, have frequently investigated the population abundance distribution. When analyzing synaptic population data, we suggest that the *shapes* of distributions (as opposed to just their means and standard deviations) have biological relevance, and may reflect a balance of the above dynamic processes operating on synapses. This is suggested both from a data-driven point of view, as skewed distributions are known to be common (Buzsáki & Mizuseki, 2014; Moser et al., 1997; Song et al., 2005) and from a functional standpoint, as large synapses have been suggested to have outsize importance (Levi et al., 2022; Omura et al., 2015; Shapson-Coe et al., 2024). In this study we aim to find a suitable low-dimensional descriptor for the shapes of synaptic populations. We then proceed to explore and contrast regions in the mouse brain, at various spatial scales, and across the lifespan of the animal, employing such a fingerprint as the main tool of analysis. We furthermore evaluate this descriptor as a means to obtain a data-driven hierarchical description of synaptic populations in the brain.

We analyze the distribution of PSD95, a key postsynaptic protein, which participates in the anchoring of AMPA and NMDA receptors and is involved in short-term and long-term plasticity of the synapse (Carlisle et al., 2008; Cuthbert et al., 2007; MacGillavry et al., 2013; Migaud et al., 1998; Schnell et al., 2002). Importantly, the level of PSD95 in a synapse has been closely linked to synaptic efficacy and stability. It displays a linear relationship to the EPSC amplitude, with a correlation coefficient of 0.8 (Melander et al., 2021). On a regional level, area measures of PSD95, of the type that we use, scale linearly with synaptic apposition face areas (Santuy et al., 2020).

## Methods

In previous work (Cizeron et al., 2020; Zhu et al., 2018) we considered *multivariate* morphological characterizations of individual synapses, and discovered that synapses exhibit large diversity. We referred to this high-dimensional universe of individual synapses as the *synaptome*. In the present study we instead described each synapse with a single *scalar* quantity, chosen to correlate well with synaptic efficacy. And while the object of study in our previous work was the individual synapse, we here shifted our attention to the collective properties of synaptic populations. We refer to sets of fingerprints of the distribution properties, which then are only defined on the population level, as the *distronome*. This description is thus complementary to the one used in Cizeron et al. (2020). As we will see, regional fingerprints based on the distronome are convergent in part with both anatomical and molecular approaches, but also deviate from those in notable ways.

Specifically, we estimated the total amount of eGFP tagged PSD95 protein in a synapse, in arbitrary units, by a measure that we call *total intensity*. Total intensity is the integral of an optical fluorescence signal over a small, algorithmically delineated area known as a *punctum*. More details on this measure can be found in the Supplementary methods section.

Complete parasagittal sections of mouse brains were delineated according to the Allen Brain Atlas (Cizeron et al., 2020). We carried out analysis on multiple anatomical levels, ranging from major areas, such as the hippocampus as a whole, to very specific sub-regions, such as the stratum oriens of the CA1. Brain sections were prepared from male C57BL/6J mice at postnatal ages of one week (1W), three months (3M), six months (6M) or eighteen months (18M), allowing us to track changes across the lifespan, though not within a single individual.

Additionally, we subdivided sections and regions into square *tiles* with a side of 100 μm. This accomplished two things. Firstly, it reduced mixing arising from spatial heterogeneity within a brain region. Secondly, it allowed us to carry out a second-order analysis of variability across tiles within a region. Therefore, most of our analysis proceeded in two steps: first we characterized the distronomical profiles of individual tiles, then aggregated across anatomical regions. In addition, we performed data-driven analysis and clustering of tile profiles, disregarding the anatomical delineation.

In order to explore characteristics of different regions of the brain, to map local heterogeneity, and to trace the changes of synaptic distribution profiles across the lifespan of the animal, we sought to compactly describe synaptic populations. As building blocks of a tentative profile descriptor we considered classical statistical moments, robust versions of the same (Kim & White, 2004), quantiles in the upper tail, and measures of distributional obesity. Additionally, we evaluated some of these for transformed versions of the data: empirical distributions normalized in some way. We further considered log-transformed data, but ultimately chose to operate in the original linear space. The standard multivariate profile descriptor that we chose, after extensive experimentation, comprised the arithmetic mean (mean_a_), the normalized width (std_n_), robust skewness (skew_r_), robust kurtosis (kurt_r_), and the spatial synaptic density. Part of the motivation for this choice was that it is somewhat canonical, comprising essentially the first four statistical moments, plus the overall synaptic density. The precise definitions of these measures, a comparison to some alternatives, and an application of these moments to electron microscopy data can be found in the Supplementary methods section.

## Results

Fig. 1a illustrates the data used in our study. It shows the computed *total intensity* of individual puncta in one brain section, and likewise for a magnified square tile located in the hippocampus. As we will see throughout this work, characterizations based on local population distributions enable a much more informative analysis compared to the single puncta data. Aggregating the raw data according to known anatomy, we found that regional profiles vary significantly. This is illustrated for five sub-regions, from different parts of the brain, in Fig. 1b. The profiles visibly differ in the location of their modes, their widths, skewness, and kurtosis. Closer inspection also reveals pronounced differences in the upper tails of the distributions (Fig. 1c).

**Figure 1.**
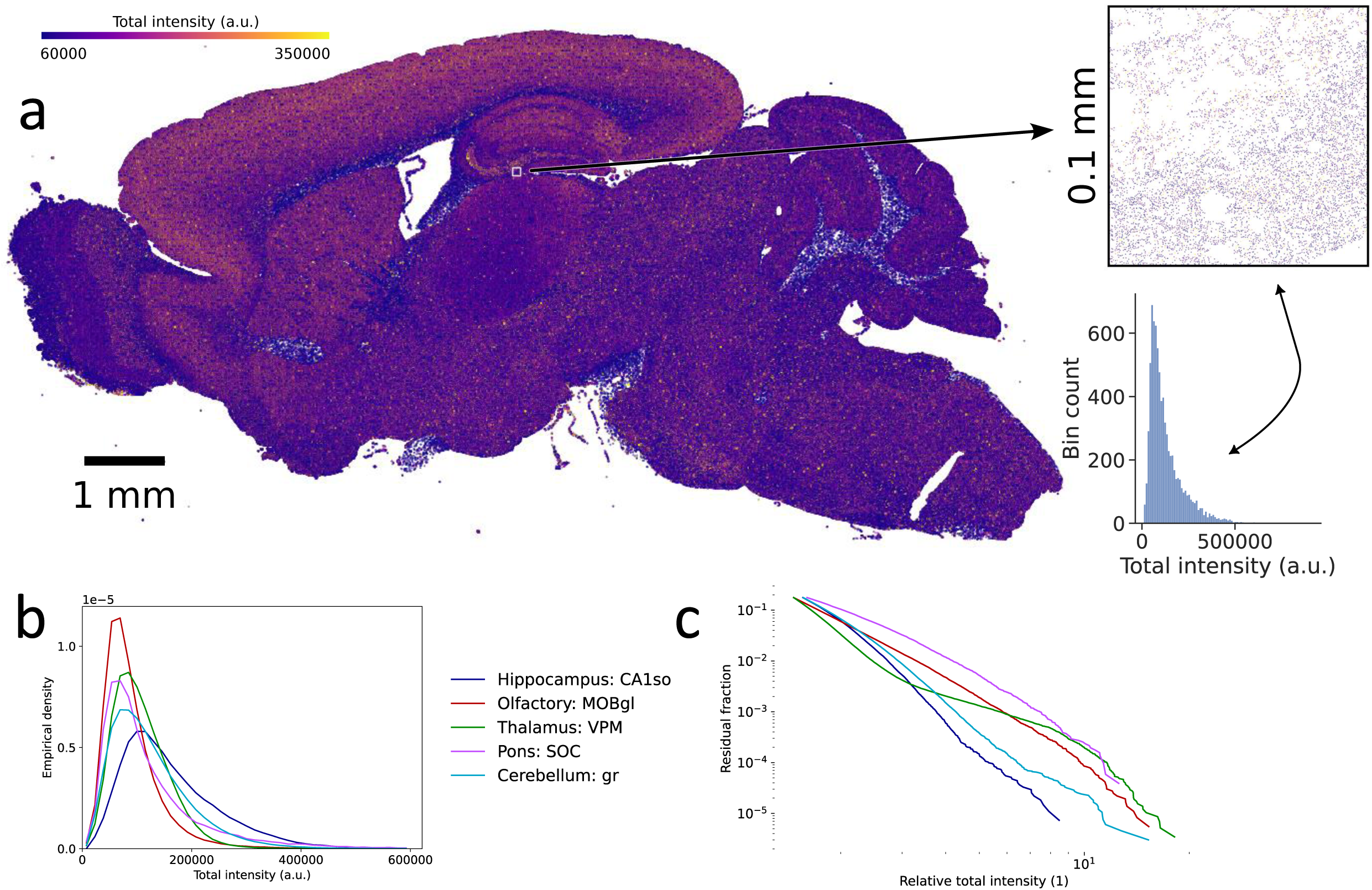
Overview of puncta data. a. A parasagittal section from a three months old mouse. Individual puncta are plotted as dots with their total intensity (see the text) mapped linearly to a color code, clipped at both ends. After filtering out extremely large and extremely dense ones, this section contains 23.9 million PSD95 puncta. The inset shows a 100 μm x 100 μm tile from the hippocampus, along with a raw histogram of the total intensity of the 7830 puncta contained therein. b. Frequency distribution of the total intensities in a few brain subregions, exemplifying the diversity of distribution shapes. In contrast to the inset histogram above, these histograms are normalized and representing an entire subregion, rather than a local tile. The regions are field CA1, stratum oriens – hippocampus, main olfactory bulb, glomerular layer – olfactory, ventral posteromedial nucleus – thalamus, superior olivary complex – pons, and granular layers – cerebellum. c. Diversity of tails. The complementary cumulative distribution function, showing the right tails (upper 15%) of the distributions from panel b, normalized to the respective distribution means.

### Regional organization of distribution types

We selected twelve top-level anatomical brain regions (cortex, hippocampus, olfactory, cortical subplate, striatum, pallidum, thalamus, hypothalamus, midbrain, cerebellum, pons, and medulla) and compared their synaptic profiles. As described in Methods and further in Supplementary methods, we parcellated the regions into tiles, profiled each tile, and analyzed the distributions of the profile descriptors over the tiles. Note, then, that two levels of “distributions” occur here, firstly the distribution of total intensities within a tile, yielding a fingerprint, secondly the distribution of tile properties within a brain region. This approach allowed us to capture distribution properties at the local level of a tile and then to characterize a region by aggregating information across tiles, in order to find e.g. the spatial variability within a region.

From Fig. 2 we can see that both the modes and spreads of all five descriptor components (mean_a_, std_n_, skew_r_, kurt_r_, and spatial density) exhibit pronounced differences between brain areas, and also strongly vary by the age of the animals. Putting questions about changes over the lifespan to one side, we first focused on the adult mice, age group 3M. We found that the upper tails of the synaptic distributions, which are reflected through the robust kurtosis measure (see Supplementary methods for a comparison to direct tail measures) spanned from relatively heavy-tailed (HT) regions, which also are more skewed, to less heavy-tailed (LT) ones, which are also less skewed (Fig. 2c-d). In general, regions belonging to the hind- and midbrain are of the HT-type, whereas forebrain regions, in particular the cortex and the hippocampus, are of the LT-type. The *extensive* measures of the synaptic distributions, the mean intensity and the spatial density, follow the opposite trend, with the highest values appearing in the anterior end, and the lowest found in posterior regions (Fig. 2a, e). In this sense, the distronome exhibits a hierarchical structure, ranging from the cortex down to the hindbrain.

**Figure 2.**
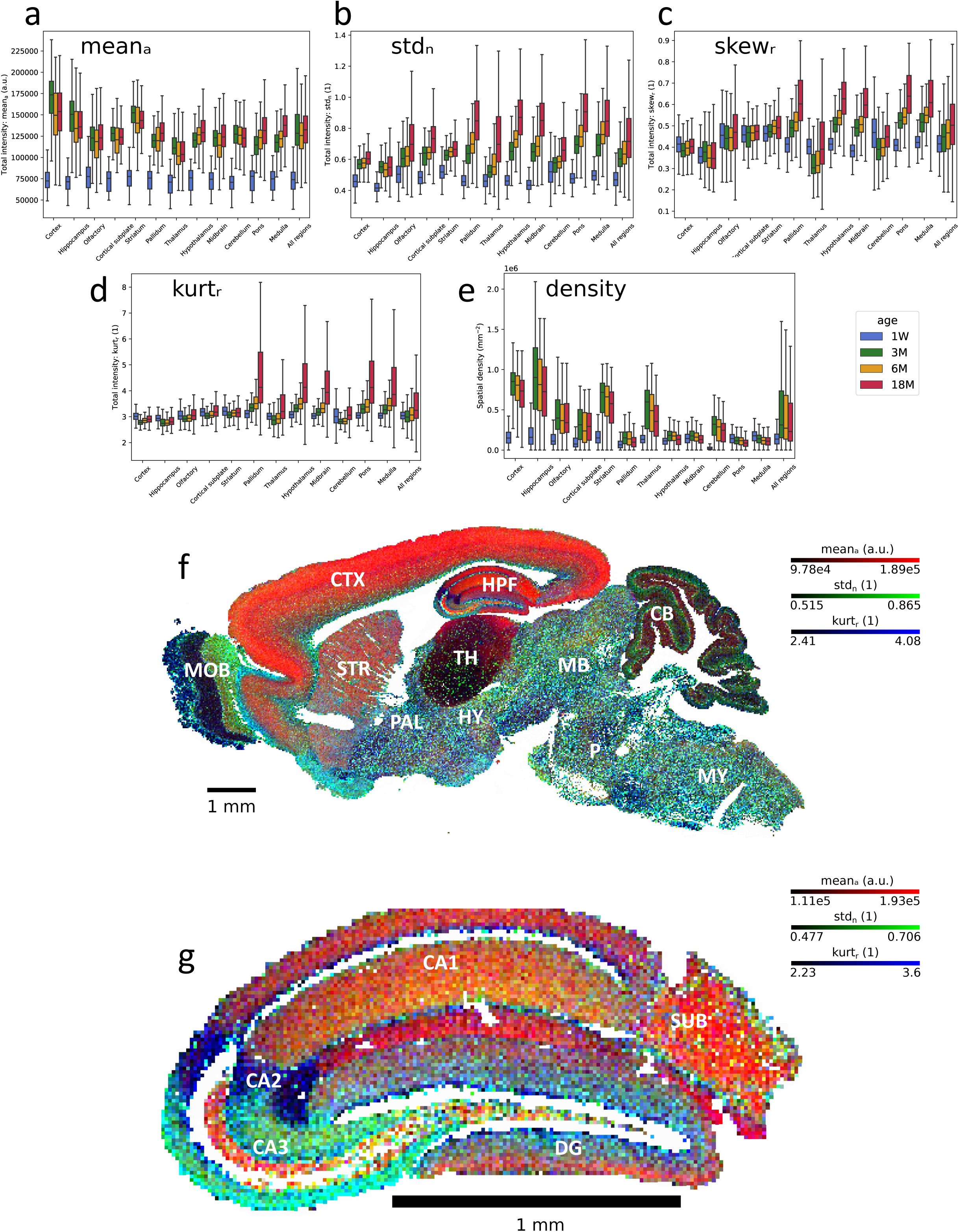
Statistical moments and dimensionality reduction. a—e. Statistical summaries of the puncta profiles for major brain regions. The statistics are computed per tile (100 µm x 100 µm), filtering out extreme puncta and excluding tiles with fewer than 50 remaining puncta. The data is aggregated by age. There were N = 5 individuals in each of the five groups 1W–18M, representing 1 week through 18 months of postnatal age. The regions are sorted from anterior to posterior, with the rightmost bar (“All regions”) the aggregate of all regions. a. Arithmetic mean. b. Standard deviation, normalized by the mean. c. Robust skewness (see Methods). d. Robust kurtosis (see Methods). e. Spatial density (puncta count per tile area). These five statistics (meanₐ, stdₙ, skewᵣ, kurtᵣ and spatial density) taken together constitute a profile descriptor that we will use to characterize regions and areas. f. Global distributional structure. False color representation of three statistical moments, in a three month old individual. Tile size 25 µm x 25 µm. The tiles are color coded by meanₐ (red), stdₙ (green) and kurtᵣ (blue), clipped at the 5th and 95th percentiles. The areas indicated are CTX: cerebral cortex, HPF: hippocampal formation, MOB: main olfactory bulb, STR: striatum, PAL: pallidum, TH: thalamus, HY: hypothalamus, MB: midbrain, CB: cerebellum, P: pons, and MY: medulla. g. Hippocampal distribution structure. False color representation as in f. Tile size 12.5 µm x 12.5 µm. Profile descriptors are coded according to meanₐ (red), stdₙ (green), kurtᵣ (blue). Tiles with fewer than 50 puncta, here mainly cell body layers, are coded white in both f and g. The areas indicated are the subfields CA1, CA2, CA3, DG: dentate gyrus, and SUB: subiculum.

We can also describe the distribution profiles, to a first approximation, as tracing the anterior-posterior neuraxis. Namely, the telencephalon is characterized by high intensity, but less heavy-tailed and skewed profiles, whereas the mesencephalon and the rhombencephalon (except the cerebellum) exhibit more heavy-tailed, low intensity ones. From the cerebellum our data set only selects the cerebellar cortex. This is characterized by low values for all five profile descriptor components and the cerebellar cortex thereby shares a similarity to the LT regions, in terms of its lower values for std_n_, skew_r_ and kurt_r_. In the diencephalon, the thalamus is similar to the cerebellum, while the hypothalamus resembles the midbrain and hindbrain regions (HT-type). The split into HT and LT regions is fully consistent with a dichotomy found using spatial transcriptomics (Yao et al., 2023) of anterior versus posterior regions. We will discuss groupings of brain regions further in the section “Hierarchical analysis and comparison to transcriptomics”.

### Homogeneity and heterogeneity

In addition to the differences in their modal profiles, we found that some major anatomical regions appear internally quite homogeneous, whereas others show much greater diversity between their constituent subregions. Some observations on this follow in this section and we will return to patterns of heterogeneity in the Discussion. We noted that pairwise comparisons between, for instance, the 12 main regions, the 4 major subdivisions of the hippocampus, as well as between its laminar subdivisions, in almost all cases reveal statistically significant differences, even when considering the individual components of the profile descriptors in isolation. The quantitative results, along with statistical tests can be found in Table 1 and Supplementary Table S1.

**Table 1.**
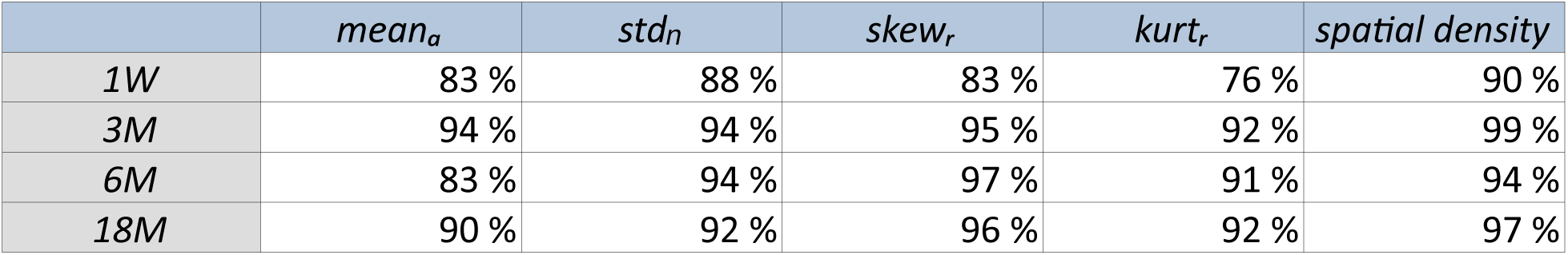
The fraction of pairs across the twelve main regions where each moment was found to differ significantly, at the p < 1% level. The comparison was done separately per age group.

As described above, a gross anterior-posterior gradient emerged in the modal profile descriptors when partitioning based on known anatomy. Specifically, we observed a decreasing trend for mean intensity and density and conversely increasing trends for std_n_, skew_r_ and kurt_r_. Without reference to anatomical delineations, a reduced three-dimensional fingerprint of tile profiles, composed of three of our five descriptor components, namely mean_a_, std_n_, and kurt_r_, was visualized as a color code (Fig. 2f-g). Major anatomical regions can be identified at a glance from this visual representation. Here, certain top-level anatomical areas, such as the hippocampus and the olfactory bulb, show very notable sub-structures, seemingly corresponding to finer anatomical details, whereas others, such as the cortex, when regarded along an anterior–posterior axis, do not. Additionally, anatomically distant areas in few cases appear similar, such as certain parts of the thalamus, which resemble the granule layer of the cerebellum, and the stratum oriens of the hippocampal subfield CA3 which resembles the olfactory tubercle. Later on, we will also see within-region diversity reflected through low-dimensional projections of the distronome.

### Homogeneity in the cortex, striatum, and in the thalamus

Along the anterior to posterior axis of the neocortex, profile descriptors vary little. The distribution shapes all the way from the motor cortex, via the somatosensory cortex, to the posterior parietal cortex are remarkably uniform over this large anatomical distance, despite known differences in lamination (e.g. absence and presence of layer 4), neuronal morphology, and functional modality (Fig. 3b1-2, 3a1, Supplementary Fig. S3I).

**Figure 3.**
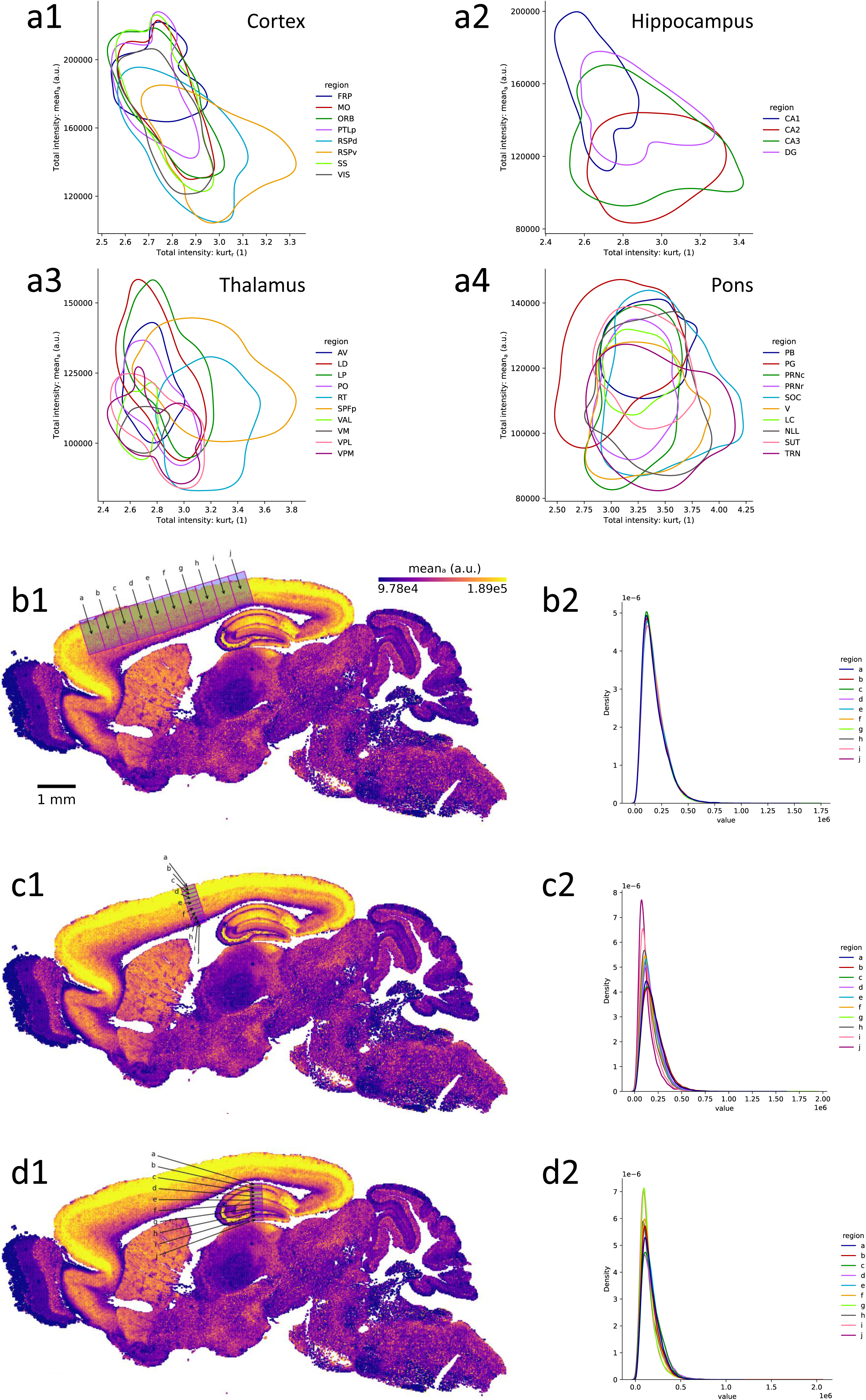
Joint structure and gradients. a. The joint distributions of two descriptor components; robust kurtosis and arithmetic mean, per region, pooled across the three months individuals. Tile size 100 μm x 100 μm. Contour plots at the 25th percentile. a1. Cortical areas. a2. Fields of the hippocampus, and the dentate gyrus. a3. Thalamic nuclei. a4. Parts of the pons. b—d. To the left (1) are the locations of ten boxes a—j in a section from a three months old individual. To the right (2) are the corresponding normalized total intensity histograms, showing data pooled across individuals of the same age, where the box locations were registered across the individuals. b. Tangential gradient (anterior to posterior) in the cortex. c. Radial gradient (superficial to deep) in the somatosensory cortex. d. Transverse gradient (dorsal to ventral) in the hippocampus.

Along the radial axis of cortex, from the superficial layer 1 to the deep layer 6, there is a minor systematic trend from wider and more light-tailed distributions to narrower and less light tailed distributions (Fig. 3c1-2, Supplementary Fig. S3II-S3V). The mean_a_ measure displays an inverted U-shape with a maximum around layer 4 (or superficial 5 for the agranular cortices) and small values particularly in layer 6. The value of std_n_ is relatively constant through layers 1-5 and increases somewhat for layer 6. The spatial density decreases, with an almost linear decrease from layers 2/3 to 5. Like the cortex overall, we found the striatum to be relatively homogeneous, except for the islands of Calleja, which have a lower mean but higher values for the other moments.

For the most part, the thalamus appeared relatively homogeneous as well (Fig. 3a3, 4c). We were nevertheless interested to see if robust differences could be found between its nuclei. Statistical analysis showed that all the thalamic nuclei could indeed be separated based on their profiles. Some profile components differ by as much as 15% when comparing an individual nucleus to the thalamus as a whole (Supplementary Table S1). Interestingly, the reticular and the subparafascicular nuclei appear to be very distinct from the rest of the thalamus, and much more similar to the midbrain and hindbrain.

### Heterogeneity in the hippocampus and in the olfactory bulb

The hippocampus shows significant divergence between the subfields CA1-3, along with the dentate gyrus (Fig. 2g, 3a2, Supplementary Fig. S1e-h, S3VI, S4), such that 84% of the pairwise single-component comparisons are significant at p<0.01. Note for instance in Fig. 2g the differences in the hues of CA1 (slm red; sr and so, orange), CA2 (blue), CA3 (turquoise), CA3 sl (orange), and DG (mixed colors). The laminar profiles per subfield also differ considerably (Supplementary Fig. S4a-d, Supplementary Table S1) where 76% of comparisons are significant at p<0.01. CA1sr, CA1slm, CA1so, and CA3slu have profiles similar to the cortex. CA2sp displays a relatively large std_n_, whereas CA2so, CA2slm, and CA3slm have small std_n_. CA2sr has a low mean_a_ and a low kurt_r_ relative to the hippocampus as a whole. In radial sections through the hippocampus (Supplementary Fig. S3VI), from CA1so, via CA1slm and the dentate gyrus molecular layer of the superior blade, to the molecular layer of the inferior blade, we particularly noted that the mode of the distribution (location of the peak) is remarkably constant, such that the radial change is mainly in the width and shape.

The olfactory bulb also displays very pronounced differences between its subparts, where the granule layer of the main olfactory bulb is light-tailed but relatively broad, while the glomerular layer is more heavy-tailed (Fig. 2f, Supplementary Fig. S3VIII).

### Within-region diversity on the tile level

Taking anatomical delineations as the starting point, our analysis above revealed that the top-level anatomical regions span a wide range along a homogeneity–heterogeneity axis. For instance, the cortex appeared very homogeneous, the hippocampus very diverse. However, this conclusion is affected by the partitioning of regions into subregions, which could be argued to be somewhat arbitrary. We were also interested in investigating this question from a more data-driven perspective and thus turned to assess diversity on the level of tiles. Returning to Fig. 2, we read off the internal heterogeneity of regions (considering single descriptor components) by looking at the spans of the bars (which indicate the 25^th^ to 75^th^ percentiles). For pairs of components, the diversities of some regions are further shown in contour plots (Fig. 3a, 4, 5, Supplementary Fig. S6, and S7).

**Figure 4.**
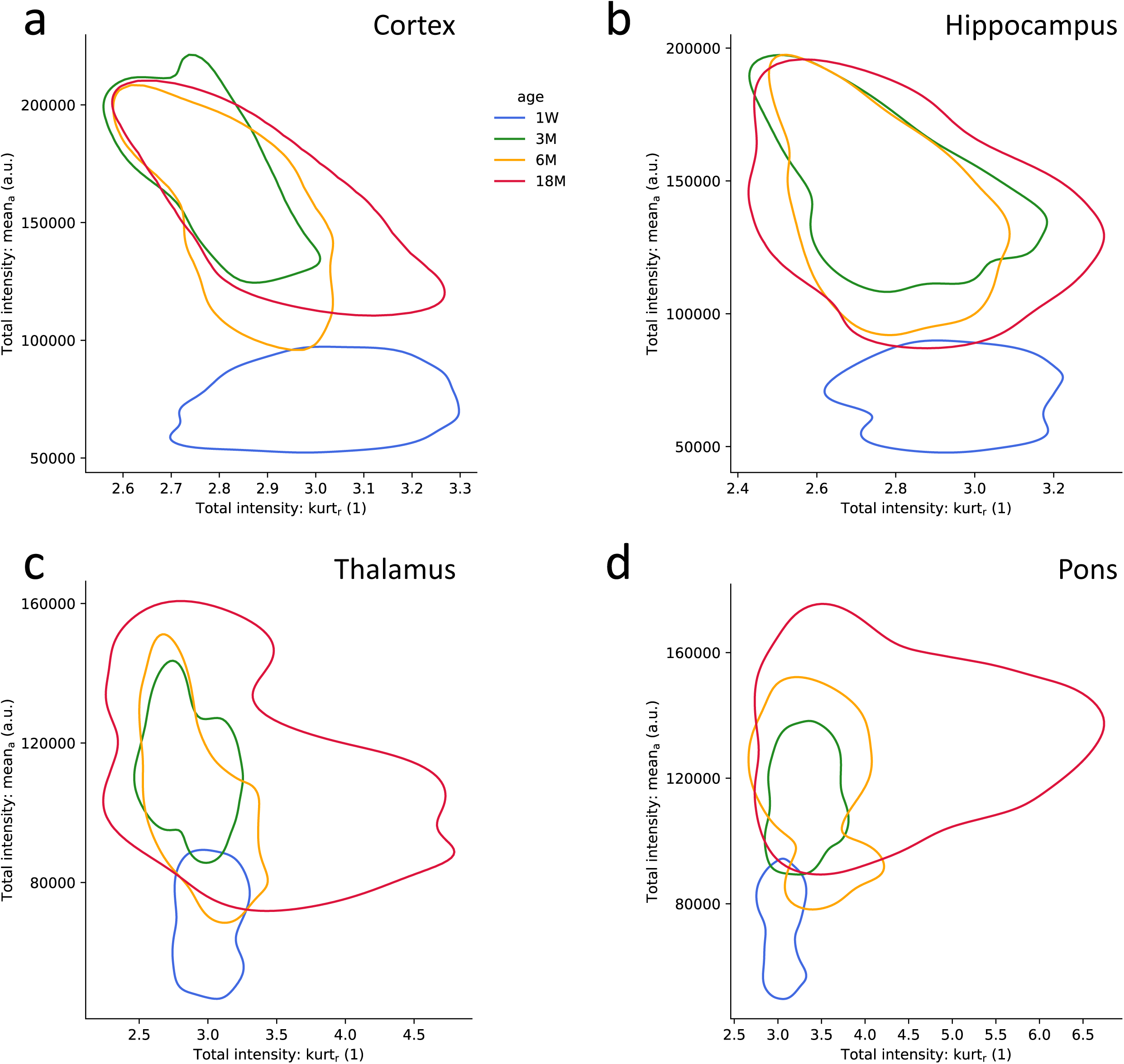
Profile evolution over the lifespan. a—d. The joint distributions of two descriptor components; robust kurtosis and arithmetic mean, per region and age group, pooled across the individuals. Tile size 100 μm x 100 μm. Contour plots at the 25th percentile. a. Cortex. b. Hippocampus. c. Thalamus. d. Pons.

**Figure 5.**
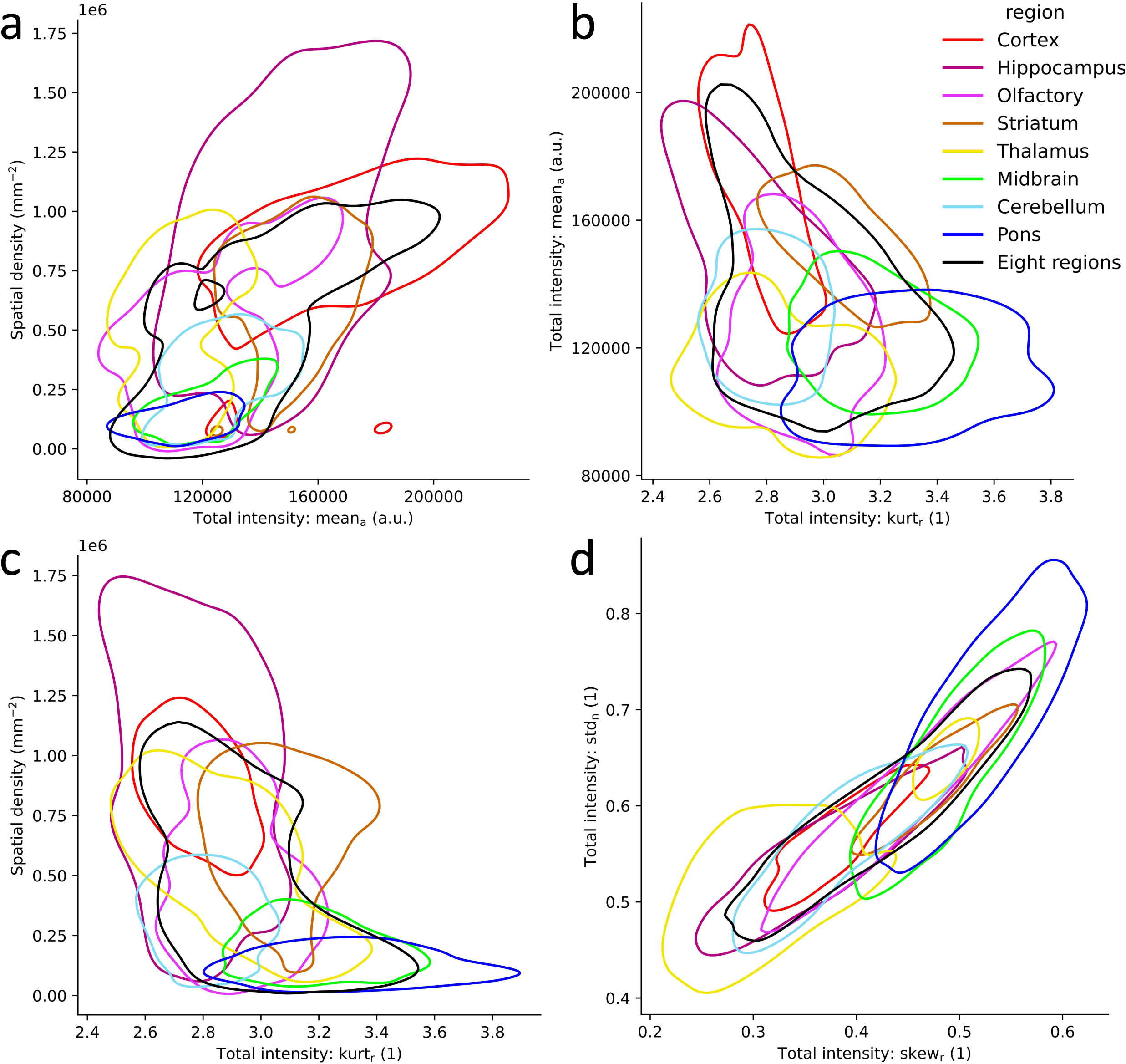
Relations between statistical moments. a—d. The joint distributions of pairs of descriptor components, per region, pooled across the three months individuals. “Eight regions” refer to the joint data across the eight regions shown. Tile size 100 μm x 100 μm. Contour plots at the 25th percentile. a. Arithmetic mean vs. spatial density. b. Robust kurtosis vs. arithmetic mean. c. Robust kurtosis vs. spatial density. d. Robust skewness vs. normalized standard deviation.

The spreads of each descriptor component per region are tabulated in the second sheet of Supplementary Table S2. We found that in the LT-regions the extensive measures (mean intensity and spatial density), which are there large, compared to in the HT-regions, vary greatly spatially, whereas the moments pertinent to shape, there small, are rather homogeneous within those regions. For the HT-regions we found a mirror image pattern. In these regions the density and the mean intensities are low and vary moderately spatially, but local variation then manifests instead through the higher spreads in skewness, and even more so in the robust kurtosis. This means in turn that in these regions the very *local* (within 100 μm) synaptic populations span a range from relatively homogeneous (smaller std_n_, skew_r_, and kurt_r_) to very diverse.

To further map the regional heterogeneity, we compared for each profile measure its standard deviation across the tiles of a region to the mean of the same. We here found a very large correlation for density and a large correlation for kurt_r_. There is modest correlation for mean_a_ and std_n_ but no apparent correlation for skew_r_ (Supplementary Fig. S5).

### Lifespan

#### Uniformity at birth

The distributions of synapses at 1 week postnatally (“1W”) stand in stark contrast to the mature brain. At that early time distributions are very similar across all regions and characterized by low mean intensity, low skewness, and very low kurtosis. The synaptic distributions resemble Gaussian distributions (Fig. 2a-e, Supplementary Table S1). In the adult at 3M, as was discussed above, regions show a pronounced regional diversification, which over the lifespan further diversifies. There is an increase in mean intensity between 1W and 3M across the entire brain , indicating that synapses acquire more PSD95, while the distribution shapes diverge. Thus, for the LT-regions, such as the cortex and the hippocampus, the tails of the distributions generally get thinner between 1W and 3M, whereas the HT-regions, such as the hypothalamus and the pons, grow fatter fails (Fig. 2d, 4, and Supplementary Fig. S6). Overall, the changes over the lifespan are strongly significant (Table 2, Supplementary Table S2).

**Table 2.** The fraction of the twelve main regions where each moment was found to differ significantly, at the p < 1% level, between the three months (3M) age group on the one hand, and the other age groups (1W, 6M, 18M) on the other.

|  | *mean<sub>a</sub>* | *std<sub>n</sub>* | *skew<sub>r</sub>* | *kurt<sub>r</sub>* | *spatial density* |
| --- | --- | --- | --- | --- | --- |
| *1W* | 100 % | 100 % | 85 % | 100 % | 100 % |
| *6M* | 92 % | 92 % | 85 % | 85 % | 77 % |
| *18M* | 77 % | 100 % | 92 % | 100 % | 92 % |

#### Divergence and persistence over adult life

As in previous work (Cizeron et al., 2020) we found that over the adult lifespan, 3M to 18M, in all of our 12 main regions (except possibly for the cortical subplate) there is a loss of synapses (Supplementary Table S3), as the spatial synaptic density decreases (Fig. 2e, Supplementary Table S2). This happens through an overall downwards shift of the distribution of densities, affecting quantiles both above and below the mode. We found that most of this synaptic loss could be modeled as being proportional to the density at 3M, but with the pallidum, the pons, and the thalamus exhibiting more pronounced drops. Considering the change from 3M to 6M the picture is a bit more complex. The cerebellum and the medulla see large drops in synaptic density, whereas the pons and the hypothalamus remain mostly unchanged.

Over the adult lifespan, we found that most LT-regions show a minor decrease in mean_a_ (Cizeron et al., 2020) (Supplementary Table S2) whereas the opposite holds true for the HT-regions, leading to an overall convergence in mean. There is an increase in std_n_, skew_r_, and kurt_r_ everywhere (except for skewness in the hippocampus). These changes to the distribution profiles are more pronounced in the HT-regions and in the thalamus. The trend towards an increase in distribution width is consistent with an increase in variability observed in aging (Grillo et al., 2013). For the cortex, we saw a generally decreasing mean_a_ (Fig. 2a, Supplementary Fig. S3I-Vc), as well as increases of std_n_, skew_r_, and kurt_r_ (Fig. 2b-d, Supplementary Fig. S3I-Vd-e). There is a decreasing trend for the density (Fig. 2e, Supplementary Fig. S3I-Vf), as in previous findings in humans, non-human primates, and rodents (Petralia et al., 2014). Furthermore, we saw a small age-dependent decrease in density for the superficial layers of the most anterior cortical regions (Supplementary Fig. S3II), consistent with earlier findings (Aguilar-Hernández et al., 2020; Wallace et al., 2007) (for Wallace, in rat) and possibly also a decrease in the superficial layers of the early sensory cortices (Supplementary Fig. S3IV-V) (Calì et al., 2018; Leuba, 1983), but see Mostany et al. (2013). Moreover, our radial profiles showed a reduced density, as well as a small increase in tail weight in the deeper cortical layers (Supplementary Fig. S3II-S3V) that correspond to what is seen in the human cingulate cortex layer III (Benavides-Piccione et al., 2013). In the striatum, we observed increases in std_n_ and skew_r_ (Table 1, Fig. 2) consistent with the changes observed by Parajuli et al. (2021), however kurt_r_ shows a small decrease in their observations (per our estimates from their raw data, gratefully acknowledged) and a small increase in ours. For the olfactory bulb, in particular its anterior subdivisions, we saw a potential increase in mean_a_ (Supplementary Fig. S3VIIIc).

We found that regions which at the adult age of 3 months were light tailed remain so, whereas regions which were heavy-tailed become progressively even more heavy-tailed (Fig. 2d, 4, Supplementary Fig. S2a-b, and S6, Supplementary Table S2). For the HT-regions, we notably observed large increases in tails from 3M to 6M, and very large ones from 6M to 18M (Fig. 2d, 4d, and Supplementary Fig. S6d, Supplementary Table S2). Among the LT-regions, the striatum shows the smallest changes overall (Supplementary Table S2), while the hippocampus stays more or less unchanged in CA2slm and DGmo (Supplementary Fig. S4f, h). In this latter group, the thalamus stands out, displaying relatively larger increases of its outer tail, so that at 18M t he outer tail in fact grows quite heavy (Supplementary Fig. S6c, f). Deeper layers of the cortex show an increase for the very outermost part of their tails (Supplementary Fig. S6a, f). We also observed that changes in regional tail heterogeneity (the distance between the central measure quantiles) correlate with the changes in tail weight, so that the HT-regions show large increases in both (Fig. 2d).

Overall, the trajectory of the LT-regions is that they acquire higher intensities going into adulthood, followed by a smaller reduction in aging, while their higher moments grow only modestly during this latter phase. This applies to the cortex, the striatum, and to the olfactory regions, while in the hippocampus, as an outlying case, we saw some regions even reduce their higher moments in the older animals. We furthermore found that all regions show continuous changes in one or several population moments also for the interval 3M to 6M (Fig. 2, Supplementary Table S2).

From 3M to 18M the cortex is losing 13% of the total number of synapses and the hippocampus 24% in our data (Supplementary Table S3), while at the same time the shapes of the distributions, as observed through their tails, are largely preserved. This meshes with clear trends in the synaptic densities, though we could not rule out an increase in the spatial density of the cortical subplate (Supplementary Table S2). In the cortex, specifically, we observed a anterior-posterior gradient of decline, where the anterior end (ORB, MO) appears unchanged and the posterior end (PTLp, VIS) decreases the most (Supplementary Fig. S3If).

In contrast, the HT-regions grow strikingly thicker tails and also increase in internal non-homogeneity as the animal ages. Thus, over the lifespan, the regulation of synaptic populations appears to behave differently across regions, leading to different outcomes in what is conserved and what changes.

#### Relations between profile measures

We were interested to further map the relations between the five descriptor components. To the extent that co-variations can be found, this may imply that distribution shapes follow particular “rules”, that could give further hints on how such patterns arise and develop in the brain. In Supplementary results we report in detail on analysis carried out on the level of local tiles, and compare the results across regions and ages (Supplementary Table S4, Fig. 5, Supplementary Fig. S6, S7). We found a between-region correlation between the spatial density of synapses and their mean total intensity, both having an inverse relationship to kurt_r_, and also to skew_r_ and std_n_. Partitioning regions according to patterns of correlations yields one group with relatively higher correlations between density and mean_a_, which largely overlaps with the HT areas, and another with higher correlations between std_n_, skew_r_ and kurt_r_, which overlaps with the LT areas. We also compared our data to reported estimates of neuronal densities (Keller et al., 2018) and found a positive correlation between the densities of neurons and synaptic densities (Supplementary Table S5, r=0.36; if hypothalamus were to be excluded, then r=0.71).

#### Single factor bilinear model

Given our observation that a reduction in the extensive descriptor components, mean_a_ and density, was accompanied along the anterior–posterior axis by increases in the pure shape measures, skew_r_ and kurt_r_ (Fig. 2), we asked how much of the variability in the regional profiles could be attributed to a single explanatory factor, and how that factor would then map to the anatomy. To this end we constructed a bilinear model and used it to arrange regions along an abstract explanatory axis (Fig. 6a, b; for details, see Supplementary methods). This model ended up explaining a large portion of the regional differences (Supplementary Fig. S8). As noted before the thalamus stands out, and indeed it falls somewhat outside of the emerging pattern.

**Figure 6.**
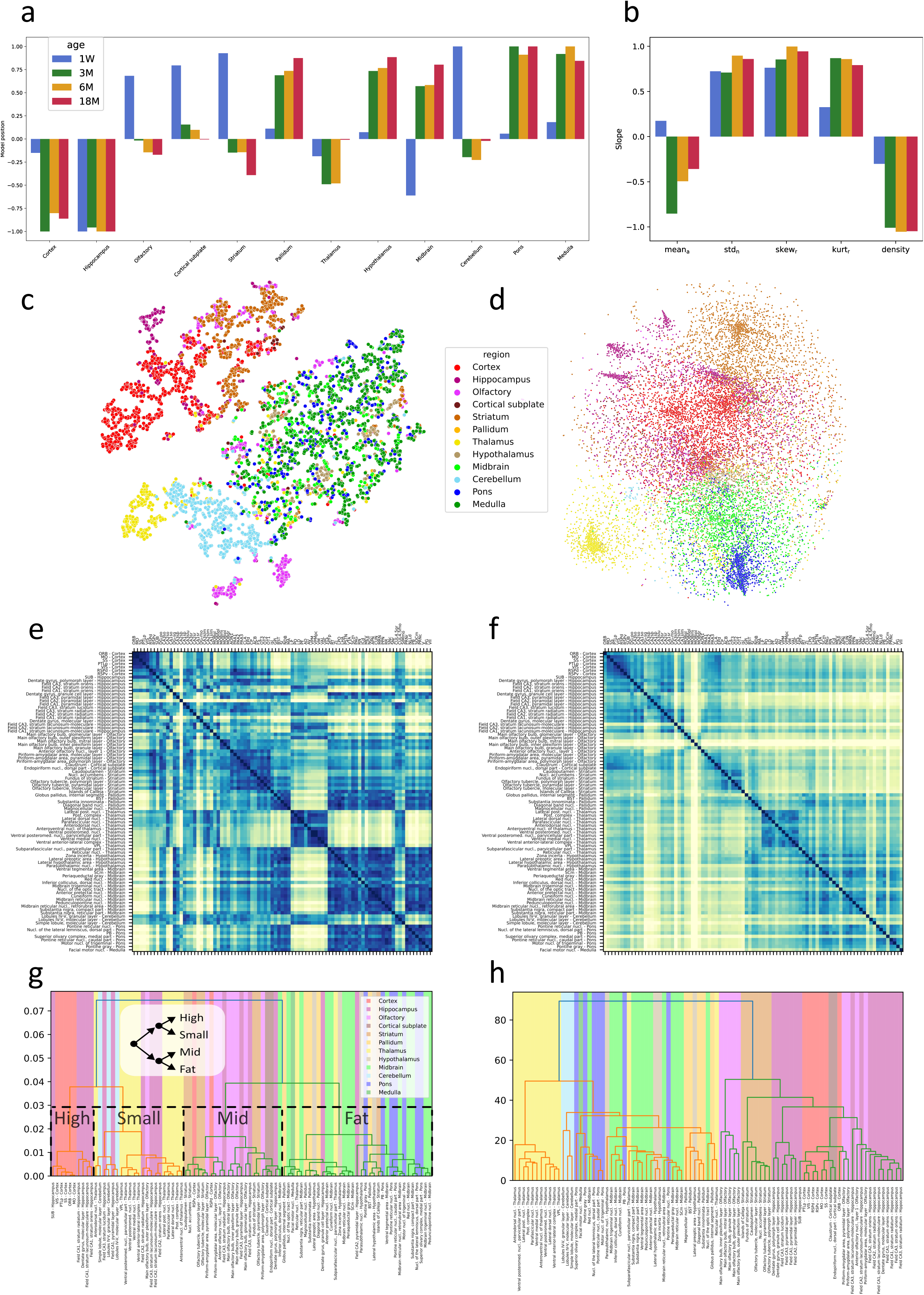
Bilinear model and comparison to gene expression data. a—b. Graded ranking of regions, using data-driven single-factor allocation. A bilinear model was built from the standardized statistical signatures of 100 µm x 100 µm tiles, with region membership as the explanatory variable. The analysis was done independently per age group, pooled across the individuals. a. Discovered regional allocations, clamped to the interval [-1, +1]. b. Regression coefficients of the regional mapping, showing the slopes of the standardized profile descriptors with relation to the regional allocation. c. Similarities between region-coded tiles. The five-dimensional profile descriptors of 100 µm x 100 µm tiles were standardized by column, and mapped onto two dimensions using a t-SNE projection. d. Similarities between spatial transcriptomics samples filtered according to the concensus synaptic gene set, and mapped onto two dimensions using a t-SNE projection. e—f. Matching the profile signatures to genetic expressions. Matrices showing the euclidean distances between 89 subregions. Low-distance (dark colored) entries indicate pairs of subregions with similar signatures. The subregions are ordered according to their anatomical location, along an anterior—posterior axis. e. Distances based on the standardized means of profile descriptors, aggregated across 100 μm x 100 μm tiles, pooled over the three months old individuals. f. Distances based on gene expression data. g—h. Hierarchical clustering of based on the preceding distance matrices, using the Ward method. g. Clustering based on distribution profiles (matrix e). Lines emphasize the outcome at the level of four clusters, where the clusters “High”, “Small”, “Mid”, and “Fat” are named based on their profile shapes, and the order is given by the centers of gravity of the clusters along an anterior—posterior axis. The inset shows the hierarchical relationship of these four clusters, which can also be read off from the full dendrogram. h. Clustering based on gene expressions (matrix f).

The outcome was consistent both over the adult age groups and between individuals within an age group. For the adult animals, the results were as follows. At one end of the emergent explanatory axis, we found regions from the telencephalon: the cortex and the hippocampus. At the other end were rhombencephalic regions, with regions from the diencephalon in between. Overall the ordering was thus mostly in line with anterior-posterior localization, but there were two exceptions: the thalamus and the cerebellum were both assigned to more “anterior” positions compared to their anatomical locations. These two areas deviate also in other respects from the other regions of the diencephalon and the rhombencephalon, respectively. We will return to this observation below, when comparing to transcriptomics data. The model thus indicates that the synaptic distributions follow an “axis” that only *partially* agrees with the anterior-posterior location, or with developmental subdivisions. Instead we may perhaps think of the emergent axis as a “hierarchical axis”, since it correlates with an evolutionary hierarchy of the brain: The cortex and the hippocampus would sit at the top of this hierarchy, the thalamus and the cerebellum in the middle, and the pons and the medulla at the bottom, which is also how they are ordered by the model.

#### Hierarchical analysis and comparison to transcriptomics

Brain organization has been described in terms of the embryological origin of cells, evolutionary homology, single cell gene expressions, macroscopic connectomics, and much more. We were interested in asking how regions and subregions would organize based on the synaptic population distribution shape. Our finding that brain regions and subregions could be segregated based on our five profile measures (Supplementary Table S1) further supports extending our analysis to multiple levels. Brain plots using three of the measures (Fig. 2f, Supplementary Fig. S6e) indeed suggested that commonalities exist between brain regions that are anatomically separated.

#### Mapping the distronome and the transcriptome

To visualize the *multi-dimensional* distronome we used the t-SNE dimensionality reduction method (Maaten & Hinton, 2008), projecting the five-dimensional profile down to two dimensions. Known course anatomy was largely recovered from this projection, as can be seen from Fig. 6c, where each projected tile has been color coded according to its anatomical designation. We note how the cortex, parts of the hippocampus, and the striatum form relatively small and distinct clusters. Conversely, we noted a very large and diverse cluster composed of tiles from midbrain and hindbrain regions. This larger heterogeneity in the midbrain and hindbrain regions, and their overlap, is consistent with observations based on spatial transcriptomics (Yao et al., 2023) where a larger number of genetically differentiated groups of cell types were observed.

Ortiz et al. (2020) collected gene expression data over the full extent of the mouse brain. Their dataset contains information on the expression of 15,326 unique genes across 34,053 sampling points and gave 181 local groups of sampling points anatomical labels. We identified 89 subregions that were common between our data set and theirs (listed in Supplementary Table S9). We also selected from their data only the set of consensus synaptic genes listed in SynaptopathyDB (Sorokina et al., 2022) comprising 3437 genes. We then carried out a second t-SNE mapping using this transcriptome data, yielding Fig. 6d.

Comparing the two t-SNE plots we found a striking similarity in terms of which regions are close neighbors. In particular, in Fig. 6c, based on our distronomical profiles, we see clusters corresponding to the cortex, the striatum, and parts of the hippocampus and the olfactory regions separately from a large grouping where tiles from the pons, the medulla, the midbrain, and the hypothalamus are intermixed. Adjacent to this we find a cerebellar cluster, two well separated olfactory clusters, and a thalamic one. In a similar vein, in Fig. 6d, based on the transcriptome, starting at the center of the figure, going upwards we find a cortical cluster and above this to the left hippocampus and to the right of this striatum. Going downwards, we find midbrain and below this pons. Going down and left, we first find the cerebellum and then the thalamus. Thus, similar brain regional organization arises when characterizing regions according to gene expression data or synaptic population shape descriptors.

#### Quantifying the comparison of regions

Having found that a projection of regions based on their distronome has a close relationship to anatomy, we wanted to further quantify the differences and similarities of regions from the viewpoint of distributional profiles. To explore this, we considered the euclidean distances between regions, based on standardized profile descriptors, and compared to the corresponding matrices derived from the spatial transcriptomics data of Ortiz et al. (2020), as per the above.

Inspecting the anatomically ordered distance matrices based on our distronome data (Fig. 6e), we found pronounced on-diagonal block structures, that well corresponded to the main anatomical parcellations. The transcriptome data exhibited a similar structure (Fig. 6f). We further studied how the correspondence between the two distance matrices depended on the choice of the distronomical descriptor, and found that it increased as we included more of its components, up to the point where we arrived at our full 5-component profile descriptor (Supplementary Fig. S9c, Supplementary Table S8). Given the generally localized nature of developmental cell lineages in the brain, it was unsurprising that the gene expression distances differed from the synaptic profile distances, in that the former preferably captured only local structure, near the diagonal. See Discussion for further explanation. The most prominent shared structure between the two matrices include the internal similarities of the cortex, the striatum, the pons, and most of the hippocampus. In the hippocampus, individual strata tend to group together, across the hippocampal subfields. For the thalamus, the matrices likewise agree on internal structure, namely that the subparafascicular and reticular nuclei stand out from the rest. On a higher anatomical level, we also noted a correspondence in the form of a larger block of similar regions comprising the hypothalamus and most of the midbrain. Taken together, this provides a strong corroboration of the notion that the shapes of synaptic distributions inform about brain organization.

#### Multi-level clustering

In order to turn the above observations on regional similarity into discrete groupings, we next carried out a hierarchical clustering, by the Ward linkage method (Müllner, 2011), also based on the distance matrices described above. The three coarsest levels of the clustering we obtained in this way (partitioning into 2, 3, and 4 clusters) are shown in Fig. 6e, g and Supplementary Table S6. A statistical characterization of these clusters is given in Supplementary Table S7. In particular, for the partitioning into 4 clusters, summarized in Supplementary Fig. S9a1-2, we found that overall the descriptor distances are statistically significant between clusters, supporting the relevance of this partitioning.

At the coarsest level, two clusters, we found that our 89 regions separated such that more light tailed and less skewed regions were found in one cluster and more heavy tailed regions in the other, where the latter also tended to have larger values for std_n_ and skew_r_. On the next level of clustering, regions with intermediate levels of these measures, including the striatum, the cortical subplate and olfactory areas, were separated from the heavy-tailed regions. Finally, at the third clustering level (four clusters) the light-tailed cluster was split in two. One cluster comprised regions with the overall smallest values of all descriptor components, mainly the thalamus and the cerebellum. The other sub-cluster had the largest overall values of the extensive measures: mean_a_ and density, but the second smallest values for the others, and included regions from the telencephalon, such as the cortex and the hippocampus (mainly CA1 and DG). The diversity of the hippocampus was again evidenced (see Supplementary Fig. S4) by its presence in all 4 clusters. The sequence of splits of the clusters and the resulting dendrogram are illustrated in Fig. 6g.

To further analyze the clusters, we performed a tail analysis. In Supplementary Fig. S9b we can observe how three of the clusters, namely ”high” (essentially telencephalon), “mid” (parts of the olfactory regions and the striatum) and “fat” (mid- and hindbrain) display a weak negative curvature, but vary in heaviness (larger slope indicates a lighter tail). However, for all clusters, and most so for the “small” cluster, which includes the thalamus and cerebellum, there is an interval of linear character. It is also interesting to note that whereas the “small” cluster starts out as the cluster with the lightest tail, it ends up with a tail as heavy as the heaviest one (“fat”) for the outermost part of the tail, probably related to RT and SPFp. This analysis found additional evidence for the notion that some regions like the hippocampus and the olfactory system are internally very diverse (already noted above in the subsection “Heterogeneity in the hippocampus and in the olfactory bulb”) leading to subregions being found in different clusters, whereas other regions are relatively homogeneous, like thalamus and hindbrain, where regions of each cluster together.

## Discussion

We characterized the synaptic populations of a region, subregion, or tile using four statistical moments and the puncta density. Using a simple bilinear statistical model, we found that most regions sorted along the anterior-posterior axis, in agreement with developmental subdivisions. In the hierarchical organization of the neuraxis, the telencephalon was characterized by high density and intensity but less heavy-tailed distributions, whereas the mesencephalon and the rhombencephalon (except for the cerebellum) exhibited more heavy-tailed distributions and lower mean intensities. Our findings of lighter tailed versus more heavy tailed regions are fully consistent with a regional dichotomy found through spatial transcriptomics (Yao et al., 2023). The intensity and density both decrease, while the tail weight increases along the anterior – posterior axis. We further found that the shapes of the synaptic total intensity distributions, as described by the statistical moments, informs on brain hierarchical organization. When more moments were included in the characterization of a subregion, a clearer organization emerged. The thalamus and the cerebellum stood out in the ordering of regions, leading us to refine our understanding from a purely anatomical ordering, to a hierarchical one. Again, this was consistent with findings by Yao et al. (2023), where these regions displayed unique protein profiles.

Moreover, we compared our data to that from spatial transcriptomics (Ortiz et al., 2020) using t-SNE projections and then found a large degree of similarity in terms of the emergent structure of regions. Comparing the underlying metric relations between regions, according to our distronome and the transcriptome, respectively, we found a large degree of similarity, but also some differences. Inspecting the distance matrices we found a pronounced on-diagonal block structure that corresponds well to main anatomical parcellations. Similarities in the gene expression profiles are mostly limited to cells of a common embryological lineage. This means that along the diagonal, blocks of similarity observed in the matrix from Ortiz et al. tended to also be found in our data, however off-diagonal blocks of similarity that we noticed were not found in the gene expression data. An interpretation of this is that similar functional goals, as reflected in the distronome, may be subserved in a convergent fashion by different proteins, operating as paralogs or analogs. Off-diagonal blocks of high similarity in our data set may thus inform about similarities across the brain in how different portions of the synaptic strengths are balanced, irrespective of the detailed genetic implementation of the plasticity mechanisms involved.

We were furthermore interested to explore the relation between the distribution shape and known properties of regions. We found over the anterior-posterior axis a difference in tail weight, where the telencephalic regions show comparably light tails and the midbrain and hindbrain regions heavier ones. Within the cortex, though, we found that the distribution shapes along its substantial anterior-posterior extent are almost identical. This is particularly surprising given differences in neuronal populations (agranular versus granular cortices), differences in cellular morphology, and in afferent projections. The radial profiles of the neocortical areas, on the other hand, show minor but consistent shifts in the distributions. Specifically, the location of the mode shifts with depth. The cortex is composed of a small number of principal neuronal types, together with interneurons arranged in a rather stereotyped layered structure. Looking at a region that is even more light-tailed, CA1 in the hippocampus, radial slices yielded remarkably constant value for the distribution peak. CA1 is dominated by a single layer of pyramidal neurons in conjunction with several interneuron types. The midbrain and the hindbrain on the other hand consist of very large numbers of neuronal types (Siletti et al., 2023). Perhaps the fatter tails that we observe in these regions are due to the larger number of cell types, each with its own location of the mode. The resulting distribution would then be a mixture, each neuronal type contributing a synaptic population light tailed in isolation. If this holds, the small but heavy tail seen in the thalamus could potentially come from its small population of inhibitory interneurons (Evangelio et al., 2018).

Our analysis highlights the differing proportions of large synapses in synaptic populations. The heavier tails in the posterior regions mean that the prevalence of large synapses is there increased. We focused our analysis at the 95^th^ percentile, but the pattern of differences between regions is similar when regarded anywhere between the 90^th^ and 99.5^th^ percentiles, with the exception that a small but heavy tail in the thalamus, which appears at 18 months, is most prominent at the extreme end of this range. The importance of large synapses has been studied experimentally (Levi et al., 2022; Omura et al., 2015; Shapson-Coe et al., 2024) as well as computationally (Omura et al., 2015). Authors argue that small populations of strong synapses can in some cases have a decisive effect on the function. It has been suggested that small spines act as “learning spines” and that larger spines are “memory spines” (Cheetham et al., 2014; Kasai et al., 2003). However, our work shows that the distributions in the hippocampus and the cortex stay very constant in the adult. One explanation for both observations could be that processes like homeostatic scaling continuously operate to maintain fixed population distributions, while allowing individual synapses to change. Moreover, it is intriguing to note that the less heavy-tailed regions are not only found in the forebrain telencephalon (the cortex, the hippocampus, the striatum) and in the diencephalon (the thalamus) but also in the hindbrain (the cerebellum). So, the maintenance of a less heavy tailed distribution is not restricted to telencephalic regions, but includes also evolutionary older structures. We also noted that the increase in tail weight in aging seen in the hindbrain regions is not directly explained by a larger loss of synapses, as the synapse loss is larger in the cortex and in the hippocampus.

We found that regions differ a great deal in their degrees of heterogeneity. We observed that the variation in spatial density within a region scales closely with the density itself. This linear scaling suggests that the variability is driven by macroscopic factors, since microscopic fluctuations alone would have resulted in a square-root-like scaling, as per a Poisson distribution. The scaling relation between a region’s mean value of each profile measure and its spread, considered over the tiles within an individual, is the strongest for density, moderate for mean intensity and robust kurtosis, but absent for skewness.

Moreover, the main regions of the midbrain and the hindbrain, and also the thalamus, have anatomical subregions which are relatively similar in their distronome. Exceptions to this are the nuclei RT and SPFp of the thalamus, which display properties similar to typical regions of the midbrain and hindbrain. Conversely, the anatomical subregions of the olfactory regions and of the hippocampus stood out as markedly divergent. Using a data-driven approach, assessing similarity between tiles, we further found that the LT-regions are spatially heterogeneous in terms of density and mean_a_, whereas the HT-regions have greater spatial variance in terms of the shape moments std_n_, skew_r_ and kurt_r_. When we compared LT-regions to HT-regions we further noted how density and intensity in the former are larger, and also vary more over the lifespan, as well as spatially (between tiles), whereas the shape moments stay very similar across both time and space. A major source of diversity is therefore the local level of density and mean intensity. One implication of the spatial variability is that experimental techniques that measure synaptic properties in spatially restricted locations may underestimate total variability. For the cortex and the hippocampus, the radial organization into layers appears to be one factor underlying spatial inhomogeneity. For HT-regions we found the mirror pattern that the shape moments are larger, and likewise exhibit the largest changes over the lifespan as well as spatially. For these regions density and mean intensity are spatially similar, such that the modes of their distributions are rather uniform across space, and the local synaptic populations instead vary greatly in skewness, and even more so in tail weight. What that means is that *locally* these HT populations may in fact either be quite homogeneous or have, for instance, long tails. The amounts of heterogeneity within the main anatomical regions was also illustrated in the dimensionality reducing projection of tile wise data. We noted then that the hindbrain regions medulla and pons are extremes in their breadth of shapes. As noted above, we speculate that one explanation for this could be a large neuronal diversity, as has been found in these regions by recent studies (Siletti et al., 2023).

In our analysis, we found significant correlations between the distribution descriptors taken between regions, to a lesser extent across the tiles of a region, and often between ages. For instance, between the main regions of our study, at 3M we found a positive correlation between the spatial density of synapses and mean_a_. This may be somewhat surprising, since one might have expected a smaller count of synapses to rather be compensated by larger sizes. Between regions, we further found positive correlations between std_n_, skew_r_ and kurt_r_. There appeared to be a negative correlation between these two major groups of descriptor components. This was more clear for density against kurtosis and to a lesser extent for mean against skewness. Within a region, across tiles, findings on correlations were rather generally weak or absent.

Using the profile descriptors to locally characterize regions, subregions or local populations, we analyzed data from animals at ages of 1 week and 3, 6, and 18 months postnatally. The distribution of synapses at 1 week is very similar across all regions and characterized by low mean intensity, low skewness, and very low kurtosis. However, in the adult at 3M, regions show a pronounced regional diversification, which then over the lifespan further greatly diversifies.

Most regions increase in total intensity from 1W to 3M, indicating that synapses acquire more PSD95. At adulthood (3M), pallial regions have changed substantially and are characterized by high density, high intensity, while the tail weights remain low. Conversely, hindbrain regions are still characterized by low density and low intensity but with an increasing tail weight. We furthermore found no support for a notion of an “adult plateau” between 3M and 6M where population distributions would be stable, but instead support for a continuous process of changes to the shapes of distributions.

Over the adult lifespan until 18 months of age, we found that most LT regions display a minor decrease in mean. As the HT regions instead increase in mean, this gives a notable convergence across the brain in this aspect. The normalized standard deviation increases everywhere, except for in the hippocampus. Finally, the pure shape moments tend to increase over the 3M–18M lifespan, but this is not everywhere the case. We found that regions which at 3 months were light tailed remain so. For example parts of the hippocampus, such as CA2slm and DGmo, appear to maintain a constant tail, whereas regions which were already strongly heavy-tailed become progressively more so. Thus, in terms of the proportion of larger synapses, the LT-regions are relatively conserved over the lifespan whereas the regions which at three months of age were more heavy-tailed change see significant increases. Over adulthood and aging, the evolutionary old but developmentally early to mature HT-regions thus undergo a progressive change towards a larger fraction of large synapses, whereas pallial regions show a pronounced conservation of their frequency distributions. The mean synaptic size also increases over the lifespan in these regions, indicating an actual growth of the synapses, rather than the fattening tails being an effect simply of synaptic pruning.

In previous work (Cizeron et al., 2020) we found a trend towards decreasing total synapse numbers and synapse density from 3M to 18M, thus correlating positively with a decrease in mean intensity. However, we here noted that the tail weights of the LT-regions are highly maintained, in contrast to HT-regions which show a continual increase in tail weight. The stability of the tail weight in the cortex and the hippocampus is particularly remarkable given a loss of synapses over age, which we also observed. In the context of synaptic losses we also noted that the product of density and mean (a perhaps somewhat crude proxy for the total synaptic efficacy in a local area) is *not* preserved over age: neither in absolute terms nor in the sense of efficacy ratios between regions.

Thus, regulation of synaptic populations is a lifelong dynamic process which may result in shifts towards larger populations of strong synapses, or instead conserve population distributions (Statman et al., 2014; Stepanyants & Escobar, 2011). This indicates that stability of the distribution, as it undergoes lifelong learning, and despite aging processes, such as a notable reduction in synapse numbers, is a characteristic property of the cortex and the hippocampus. However, this is not a necessary dynamical fate, as regions of the hindbrain show considerable change, despite playing essential roles in basic physiological function. One possible interpretation is that the statistics of the world, learned by the hippocampus and the cortex does not change, such that these areas need to stay “forever young” to remain in tune, whereas this driving force is not present in organ systems subserving internal physiological roles. Increases in the span of quantiles of synapses within a population may then reflect compensatory mechanisms to make up for losses in function across the organ system.

## Acknowledgments

We thank Masato Koike for sharing synaptic electron microscopy data of the striatum, Javier DeFelipe and Angel Merchán Pérez for sharing synaptic electron microscopy data of the hippocampus and cortex and Edita Bulovaite and Noboru Komiyama for discussions and comments on the manuscript. Work in the Fransén laboratory is supported by the following grants, Swedish VR-2022-01079, KTH DigitalFutures and computational resource allocations from SNIC. Work in the SGNG laboratory is funded by the Wellcome Trust (302077/Z/23/Z); the European Research Council (ERC) under the European Union’s Horizon 2020 Research and Innovation Programme (885069 SYNAPTOME); and Simons Initiative for the Developing Brain (SIDB) under the Simons Foundation for Autism Research Initiative (529085). For the purpose of open access, the author has applied a CC-BY-NC public copyright license to any Author Accepted Manuscript version arising from this submission.

## Supplementary methods

### Properties of the fluorescence microscopy data

The experimental protocol and data processing pipeline underlying this work was presented in Cizeron et al., (2020). In the present study we used the data for postnatal ages of 1 week and 3, 6 and 18 months (N=5 in each age group). We approximated the total amount of the protein PSD95 per synapse, in arbitrary units, by an estimator that we refer to as the *total intensity*. This is the integral of an optical fluorescence signal over a small, algorithmically delineated area known as a *punctum* (see Zhu et al. (2018) and Cizeron et al., (2020) for details on punctum delineation). Total intensity is then the product of mean intensity and size data as presented in the 2020 study. The computational pipeline for calculating these measures was calibrated against manual identification of puncta. We here excluded puncta with a size exceeding 80 or a mean exceeding 30000 in order to omit an extremely small fraction of objects we felt were outside the range of applicability. In Santuy et al. (2020), synaptic data acquired by the fluorescence microscopy methodology was compared to spine volume and synaptic face areas, as estimated through serial electron microscopy. The authors discuss the differences between estimates obtained using these two experimental methodologies, in particular potential under or overestimation issues of fluorescence microscopy, including omissions of very small synapses or underestimation of the size of the largest synapses.

We were interested in assessing the impact of the two methodologies, fluorescence microscopy and electron microscopy, on the statistical moments std_n_, skew_r_ and kurt_r_ used in our study (knowing that density was already controlled for and that mean_a_ is expressed in arbitrary units). To this end, we obtained, and gratefully acknowledge, electron microscopy data from the lab of dr. Koike (Parajuli et al., 2020). Their data pertain to the striatum, a region which exhibits intermediate levels of puncta intensity. We compared this electron microscopy data for ages 3M (N=177) and 22M (N=156) to our data for 3M and 18M, respectively. Using either face area or volume from their data we found, for both age groups, estimates of std_n_, skew_r_ and kurt_r_ which were all slightly larger than those from our study (about a factor of 1.4 greater for the former two and 1.2 for kurt_r_). Given that the striatum has a light tailed distribution, we can arrive at a quantitative mapping between dr. Koike’s data and ours. Namely, if we assume a lognormal distribution of the data, then there is a single shape parameter, the standard deviation (std). Starting from our data, we would need to scale this parameter by a factor 1.25, 1.55, and 1.65 for std_n_, skew_r_ and kurt_r_, respectively, to transform from our observations to those of the electron microscopy data. Importantly, we thus did not see evidence of an overestimation of the higher statistical moments in our analysis.

### Parcellation of the data

To perform data-driven analysis, we partitioned the brain data into square tiles. We chose a size of 100 μm x 100 μm, based on extensive experimentation to find a good trade off, between on the one hand achieving high spatial resolution and minimizing the mixing of data from neighboring locales, and on the other hand obtaining large-enough sample sizes for statistical analysis within the tiles. In order to settle this trade off, we looked at the variability of outcomes between differently translated tile grids, individuals in the same age group, and within known anatomical areas, all three of which should yield stable results. For the majority our analysis, a smaller tile size would in fact have been permissible, but for estimates of tail weights (see more below) in regions of low puncta density, the relatively large size of 100 μm x 100 μm was found to give more consistent results. Compare to the figure below. Note that when we partitioned anatomical regions into tiles, we first selected puncta based on the region, then carried out the parcellation, potentially yielding geometrically shifted grids of tiles between regions, but with no puncta shared between regions.

When data from tiles were analyzed in terms of statistical moments (see below), we omitted tiles with a count below 50, in order to have sufficient data for the estimation. In figures showing the anatomical location of tiles, tiles omitted this way were colored as background, in white.

We additionally developed a tool to enable the manual definition of regions of interest, either as rectangles or circular discs. Rectangles can be split into column-wise or row-wise stripes or into subtiles, which we used in our study of various anatomical gradients.

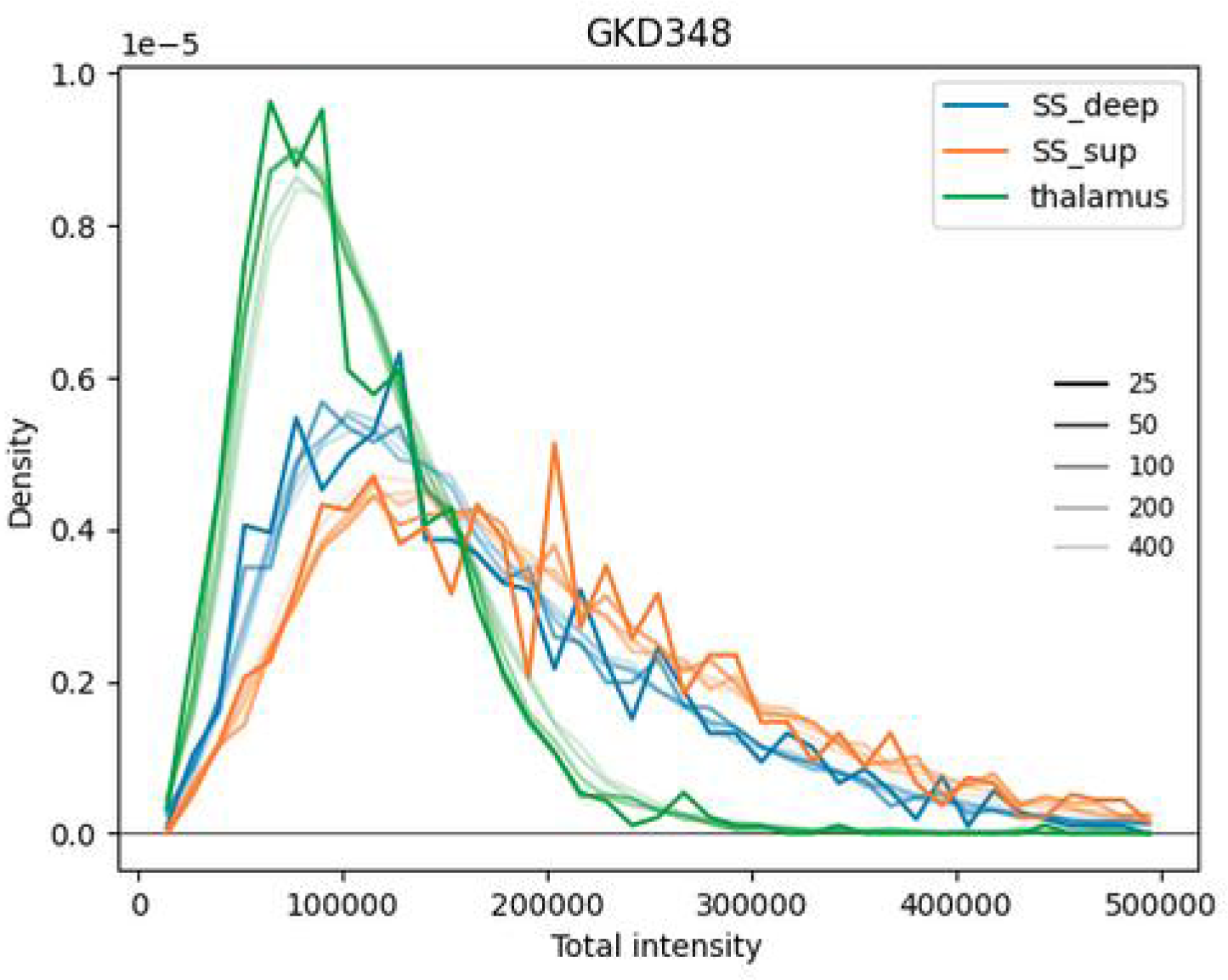

**Figure**: Frequency histogram of density of puncta vs. total intensity for tiles of different sizes. The size of the tile (side 25, 50, 100, 200, 400 μm) is shown by color intensity, and the hue indicates the brain area (superficial and deep layers of the somatosensory cortex as well as the thalamus). Data is shown for one individual from the 3M age group (ID: GKD348). Note the increasing variance across adjacent histogram bins for smaller tiles, while the overall envelope is conserved.

### Choice of moments

During initial analysis of the data, we found a strong correlation between the mean of the total intensity (mean_a_) and its standard deviation (*r*^2^ = 0.87). We therefore chose to study the relative spread instead of the variation itself, by normalizing the standard deviation by the local arithmetic mean, yielding std_n_.

We found that the classical higher statistical moments; skewness, and kurtosis even more so, were strongly dominated by the extreme tails of the puncta distributions. By definition we have limited number of observations in the tail. Scarcity of observations then translates into variability of the estimates of higher moments. We therefore used robust versions of skewness and kurtosis. This becomes particularly important for the analysis over tiles and for regions with low puncta density. Multiple robust estimators of skewness and kurtosis have been proposed in the literature. Based on Kim & White (2004) we used

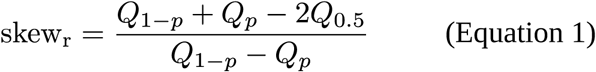

for robust skewness and

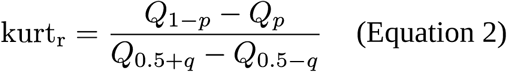

for robust kurtosis, where *Q_x_* ≡ F^-1^(*x*) are empirical quantiles, and we chose the parameters *p* = 0.02 and *q* = 0.25. The choice of *p* determines how far out on the tails the measures are evaluated, and was based on the amount of available data.

In addition, since our main interest lies in the right (upper) tails of the puncta distributions, we considered a family of upper tail descriptors, relating the inverse survival function to the central with of the distribution:

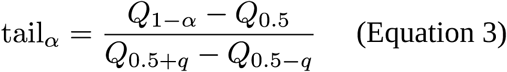

We used *α* ∈ {0.05,0.005,0.0005}, and *q* as above. We report these using percentage notation, e.g. writing tail_5%_ for *α* = 0.05.

### Statistics of regional differences

We built regression models to compare the distribution profiles between regions and subregions to each other, and to those of the brain as a whole. Because of the large differences between the age groups these models were separate by age. To handle the also notable variability between individuals within an age group, we included the identity of the animal as a categorical variable (as a set of indicator variables) in the model, thus eliminating between-individual variability common between regions and increasing the power of the model (Supplementary Table S1). From these models we were able to statistically ascertain the significance of moment differences between regions, and also to study how those differences evolve over the lifespan of the animal. Separately, to directly determine when regional profiles change over the lifespan we applied a one-way f test to the differences between observed moment values. Here we pooled the individuals within an age group (Supplementary Table S2). Because of the between-individual variability this comparison had lower statistical power than the previous model.

### Estimation of a bilinear model

In order to explore to which extent a single explanatory variable may capture the variation across brain areas, as expressed through the moments descriptor, we created a bilinear regression model.

The model is *V_jni_* ≈ *a_j_k_i_ + m_i_* where *V_jn_* = [mean*_ajn_*, std*_nĵn_*, skew*_rjn_*, kurt*_rjn_*, dens*_jn_*] is the moments descriptor vector for tile *n* ∈ [l-*N_j_*], belonging to region *j* ∈ [1..12]. Here *a_j_* can be thought of as an abstract position of the region ĵ, along an explanatory axis. We constrained this variable to lie within the interval *a_j_* ∈ [— 1,1]. Then the slopes *k_i_* specify how moment *i* ∈ [1..5] varies along this axis, and *mi* is the intercept per moment.

To fit the model, we considered the different age classes separately, and computed the moments descriptors for tilings of each main region. The individual descriptors were standardized across all tiles from the age class, to zero mean, and standard deviation one. Optimizing over *a_j_, k_i_* and *m_i_*, while observing the constraints on *at,* the tile-wise sum of squares error of the model was minimized. Finally, we expanded the explanatory axes, if needed, to span the entire interval [—1, 1], and undid the standardization of the descriptors, to report the results in their original space.

### Estimation of a hierarchical model

We build a clustering model that group brain regions based on the proximity of signatures. To capture the underlying signature of a brain region, we computed a feature vector for each of its constituent tiles. The feature vector choice that we focused on is our standard moments descriptor, though we investigated other choices as well. We then averaged the feature vectors over the region tiles (possibly across multiple animals). We finally standardized the averaged components to unit variance and zero mean, such that each one carries equal weight in the following. Next, we computed a distance matrix comprising the distances between the standardized feature vectors for all pairs of regions. The metric we choose was the Euclidean distance; another choice that we considered, leading to mostly similar results, was the cosine distance.

At this point we compared such distance matrices, calculated from the distribution shapes, to a distance matrix based on gene expression data. To this end we re-scaled both matrices to unity in the Frobenius norm, then computed the associated distance metric (the element-wise Euclidean distance) between the two matrices. These distances let us compare the overall match between the two perspectives. We also averaged distances along the rows (or, equivalently, the columns) of the matrices, to assess how well the two views match from the perspective of each region. We compared these outcomes for a few different choices of feature vectors.

Finally, we applied a greedy hierarchical clustering method, where starting from each region being its own cluster, clusters are repeatedly joined according to the Ward variance minimization criterium. We studied the clusters arising towards the end of this process, when 2–6 clusters remain. Our focus in the main text is on the level of four clusters, which we label “High”, “Small”, “Mid”, and “Fat” based on their feature characteristics.

Over a range of alternative ways to perform the clustering (different choices of features and metrics) we noted that while the clustering is mostly stable, there are some regions, such as the cortical subplate, the striatum and the olfactory regions, which may cluster differently depending on the precise choice of method.

## Supplementary results

### Relations between profile descriptors

Our analysis indicated that we could identify two main groups of brain regions, LT and HT, with distinct differences in their shape parameters and changes across the lifespan. The spatial density and the mean_a_ are large in the first group (LT) and small in the second (HT), while the converse holds true for std_n_, skew_r_ and kurt_r_. We were interested to further map the relations between statistical moments. In the following, we report on analysis carried out on the level of local tiles, and compare the results across regions and ages (Supplementary Table S4, Fig. 5, Supplementary Fig. S6, S7). To the extent that co-variations were found, this may imply that distribution shapes follow particular “rules”, that could give further hints to how such patterns arise and develop in the brain.

The correlations between the five descriptors are in general relatively weak, implying that each conveys a somewhat but not entirely orthogonal view of the synaptic distribution profiles (Supplementary Table S4). Across tiles, at 3M we found a positive correlation between the means of the spatial density and mean_a_ (r=0.34 for 3M, exemplified in Fig. 5A, 18M in Supplementary Fig. S7a2). There is further a negative correlation between on the one hand density and on the other std_n_ (r=-0.21), skew_r_ (r=-0.23), and kurt_r_ (r=-0.25, 3M exemplified in Fig. 5C, 18M in Supplementary Fig. S7a1), such that locally high density regions can be thought of as being less diverse. Meanwhile, std_n_, skew_r_ and kurt_r_ are all positively correlated (r=0.23–0.28 for 3M, exemplified in Fig. 5D, 18M in Supplementary Fig. S7a4). These relations are, despite how very different the distributions appear in the young animals compared to any adult age, somewhat present already at 1W and do persist over the adult lifespan, though at 18M we found that the correlation between density and mean_a_ was reduced. The above relationships held true also in the extreme cases. Namely, the highest synaptic densities coincided with the highest mean_a_ and with the smallest tail weights, and were found in the telencephalon, in particular in the superficial layers of the cortex (Fig. 5A-C, Supplementary Fig. S3II-V, S7a1-3), and in the hippocampal subfield CA1 (Supplementary Fig. S7c1-2). Likewise, the lowest mean_a_ and the lowest densities coincided with the largest tail weights, in the midbrain and certain hindbrain regions (exemplified in Fig. 5, Supplementary Fig. S7e). Some areas did break these rules though, such as the thalamus and cerebellum, that stood out with a combination of low mean_a_ along with low std_n_ and skew_r_ (Fig. 5a-c, Supplementary Fig. S7d).

The link between density and intensity is mostly due to between-region differences. When we considered only differences within a region, the two were in most cases not clearly correlated, though with some exceptions such as the cortex. Density and kurt_r_, and to lesser extents density and skew_r_ or std_n_ (not shown), remained inversely correlated, seen for instance in the thalamus (Fig. 5, Supplementary Fig. S7a1, S7d1). Furthermore, using reported estimates of neural densities (Keller et al., 2018), we found a positive correlation between the densities of neurons and synaptic densities (Supplementary Table S5, r=0.36; if hypothalamus were to be excluded then r=0.71).

### Summary of observations

A distronomical summary of the properties of synaptic populations constitutes a higher order signature, which can be used to characterize brain regions and subregions. By selecting a set of statistical moments, we constructed such a “fingerprint” and used it to characterize populations, mapping variations within and between brain regions. We showed that hierarchical clustering based on such a distronomical profile is consistent with hierarchical parcellations based on anatomical and on gene expression data.

Overall, we found that the shape variables, based on standard deviation, skewness and kurtosis, differentiate regions in similar ways, but distinct from the mean and the density. We further found that the telencephalic regions display fewer very large synapses relative to the midbrain and hindbrain regions which are characterized by heavy tails. The thalamus and the cerebellum are separate from both of these groups, when considering their multivariate signatures. Comparing tiles within regions to assess their spatial heterogeneity, the moments with the largest values also vary the most spatially between tiles. Tile-wise analysis shows positive correlations between standard deviation, skewness and kurtosis both in the within-region and between-region analyses and correlations between density and mean intensity in the between-region analysis.

From a common distribution at 1W of age, regions which then acquire higher intensities appear to be under tight regulatory control, whereas the regions which only increase slightly in intensity continue to change their higher statistical moments over the lifespan. In terms of the prevalence of larger synapses, the lighter-tailed regions are relatively conserved over the life-span whereas the regions which at 3M age were more heavy-tailed progressively become even more so. Moreover, whereas the higher statistical moments generally increase, there are also exceptions to this trend. Thus, over the lifespan, synaptic population regulation appears to be under different dynamic controls, which is reflected in all the statistical moments that we observed.

As in previous work (Cizeron et al., 2020), we found a trend of decreasing synapse numbers from 3M to 18M, along with a decrease in the mean_a_ intensity. The stability of the tail weight of the cortex and the hippocampus is then particularly remarkable given the loss of synapses over age, where the cortex is losing 13% and the hippocampus 24% whereas the medulla, which shows dramatic increases in tail weight does not show a clear trend. We also noted that neither the olfactory bulb nor the midbrain show any significant losses, though the range of the density decreases somewhat. In the context of synaptic losses, we also noted that the product of density and mean (a perhaps somewhat crude proxy for the total synaptic efficacy in a local area) is *not* preserved over age: neither in absolute terms nor in the sense of efficacy ratios between regions.

The distribution of properties of a synaptic population is the result of an interplay between multiple underlying functional processes, such as development, homeostasis, learning and redundancy preservation. These processes are active to variable extents over an individual’s lifetime. We propose that profiling by statistical descriptors can be powerful tools to characterize the outcomes of the underlying microdynamics, which can be hard to observe experimentally in vivo. We also propose, based on the outcomes of our analysis, that the shapes of the distributions of local synaptic populations informs on a hierarchical organizational structure of the brain.

### Supplementary tables – captions

*The supplementary tables referenced below are provided separately in spreadsheet format. Note that some tables contain multiple worksheets*.

### Cell background color codes

white = p < 0.01

yellow = 0.01 < p < 0.05

red = 0.05 < p

green = n/a

**Supplementary Table S1**

Fit of a linear regression model to quantify the difference in each of the five moments between pairs of regions. This is performed 1) at the top level 2) for the main hippocampal subdivisions 3) for subregions of the hippocampus. The models are on data from 100 µm x 100 µm tiles, pooled across the individuals within an age group. Modeling is done separately by age group. Each of the three worksheets summarize (starting from column AN) the “slope” regression parameter expressing the difference between the pairs of regions, along with its confidence interval and a significance value. Details of the underlying linear regression model are found in the columns starting from E. Columns E—H, AQ, BI, and BQ relate to the arithmetic mean, columns I—L, AU, BJ, and BR to the normalized standard deviation, and so on, for the robust skewness, robust kurtosis, and spatial density. Of these, for the arithmetic mean, the columns E —G relate to the difference in moment between the pair of regions, column E being the central estimate, columns F—G a confidence interval, and column H the p-value relative the null hypothesis. These are all summarized in column AQ, with the background color determined by the p value. Finally column BI indicates whether the p-value is below the 1% level and column BQ counts the frequency of such p-values. For the other four moments the structure is equivalent. An overall count of significant differences is given in column BV.

**Supplementary Table S2**

Means (worksheet “means”) and stds (worksheet “stds”) for each of the five moments by tile by top level region and age (like Fig. 2). For the means, color codes indicate the significance of the deviation from 3M for the other age groups. This is based on a one-way f test between the tile-wise data points for the region, pooled across the individuals within an age group. Significances between other pairs of ages can be found in the “sig_x” tabs. The worksheets labeled sig_x, where x is the name of a moment, additionally contain the f statistics and corresponding p values for the tests which make up the basis for the color code in the worksheet “means”.

**Supplementary Table S3**

Total puncta counts (mean per animal) per main region and age. Also the relative change from one age to the next one, and the count relative to 3M for the other ages.

**Supplementary Table S4**

Tile-wise correlations and also R2 values for a linear model explaining one moment in terms of another for all pairs of moments across the whole brain (but only region-tagged), per age.

**Supplementary Table S5**

The means and stds of the density per tile (like in Supplementary Table S5), by regions where we have a literature reference for cell bodies at 2M, compared to that reference value. Only 3M from our data.

**Supplementary Table S6**

Which cluster each area (in the fine spot designation of areas) falls in: High, Small, Mid, or Fat. Arranged by the coarse spot designation. Also the hierarchical structure (clustering sequence) of the four clusters.

**Supplementary Table S7**

The gridwise means (worksheet “means”) and stds (worksheet “stds”) of five moments in the four clusters. The grid-based significance (f-score numerically, p value by color [not 1:1 by f score]) of the pairwise differences of the five moments between the clusters plus between a cluster and all data within an age group. Columns E-I in the “pairwise” worksheet show f scores between the means, and color codes indicate which differences are significant, drawing on the p values in columns M-Q.

**Supplementary Table S8**

The deviation between region differences (metric) according to gene data on the one hand and moment fingerprints on the other hand. By age and for all ages. Evaluated for 1-5 moments in the fingerprints, starting with just including the arithmetic mean, then adding, sequentially, the normalized standard deviation, the robust skewness, the robust kurtosis, and finally the spatial density.

**Supplementary Table S9**

Axis text in Fig. 6e-h, Supplementary Fig. S9d1-d2.

**Supplementary Table S10**

Abbreviations used in Fig. 3 and Supplementary Fig. S7.

**Supplementary Table S11**

Abbreviations used in Supplementary Fig. S7.

---

**Supplementary Figure S1.**
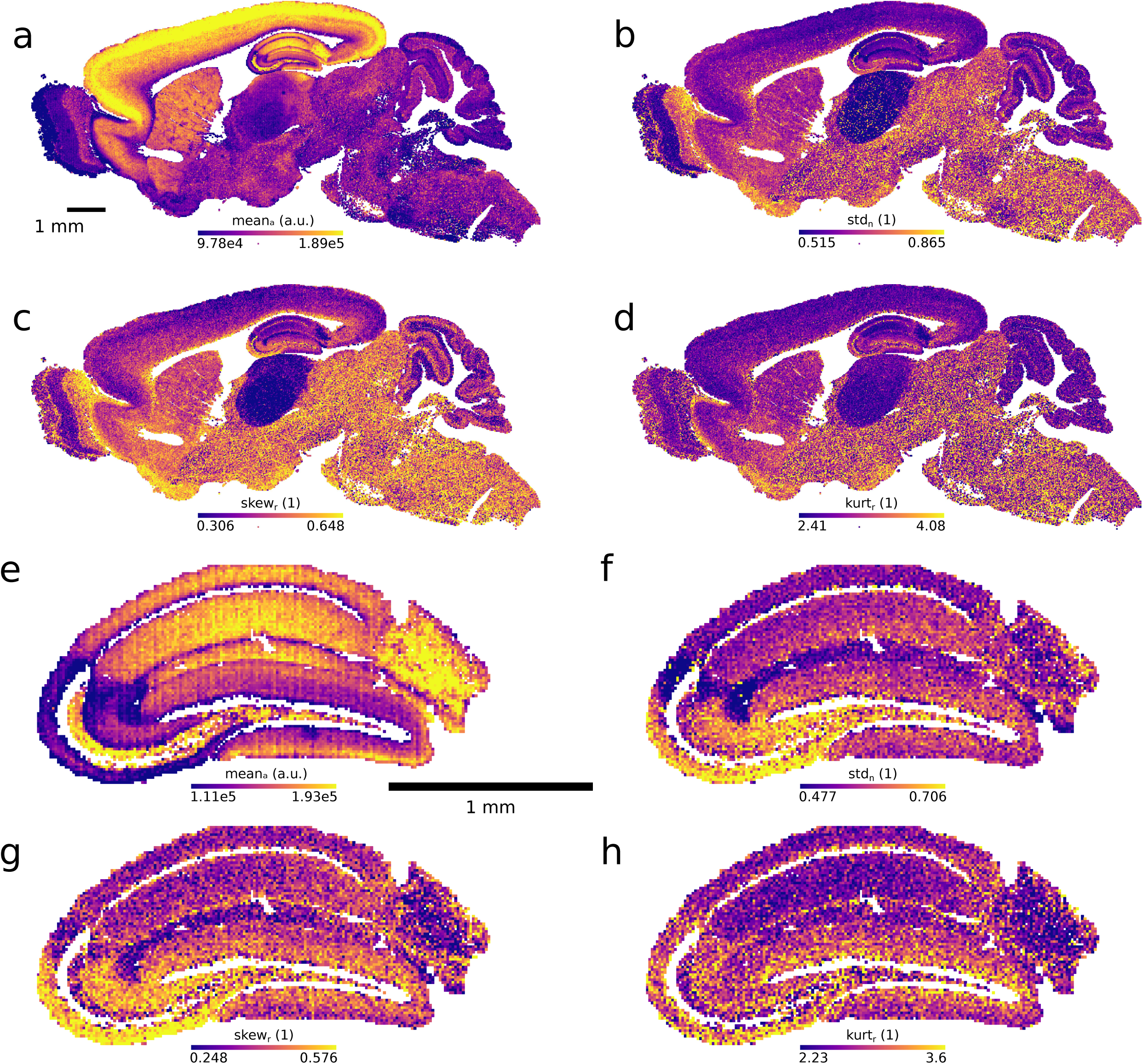
Moments overview. a—d. Full section from a three months old individual. Each tile is color coded according to the value of one moment. The color scale is clipped at the 5th and 95th percentiles. Tile size 25 µm x 25 µm. e—f. The hippocampus from the same individual. Tile size 12.5 µm x 12.5 µm. a, e. Arithmetic mean. b, f. Standard deviation, normalized by the mean. c, g. Robust skewness. d, h.

**Supplementary Figure S2.**
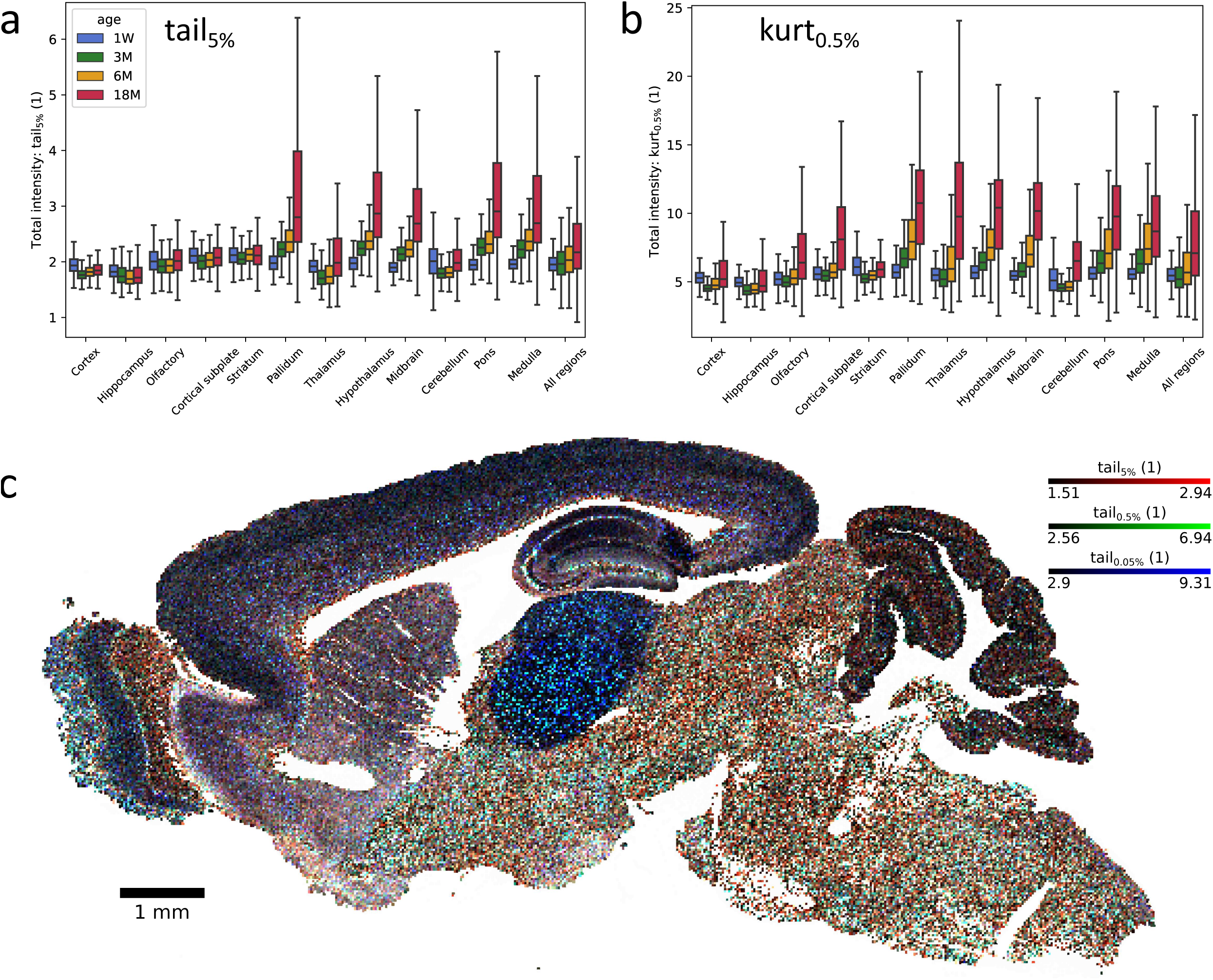
Additional statistical moments (refer to Fig. 2). a. Relative 95th percentile value (see Methods). b. Robust kurtosis at the 0.5th percentile (see Methods). c. Brain overview for a three months old individual. Each 100 µm x 100 µm tile is color coded according to the relative tail values at the upper 0.5th (red), 0.05th (green) and 0.005th (blue) percentiles (see Methods). Note how the tail of thalamus is prominent at the outermost tail value (dominated by blue hues).

**Supplementary Figure S3I.**
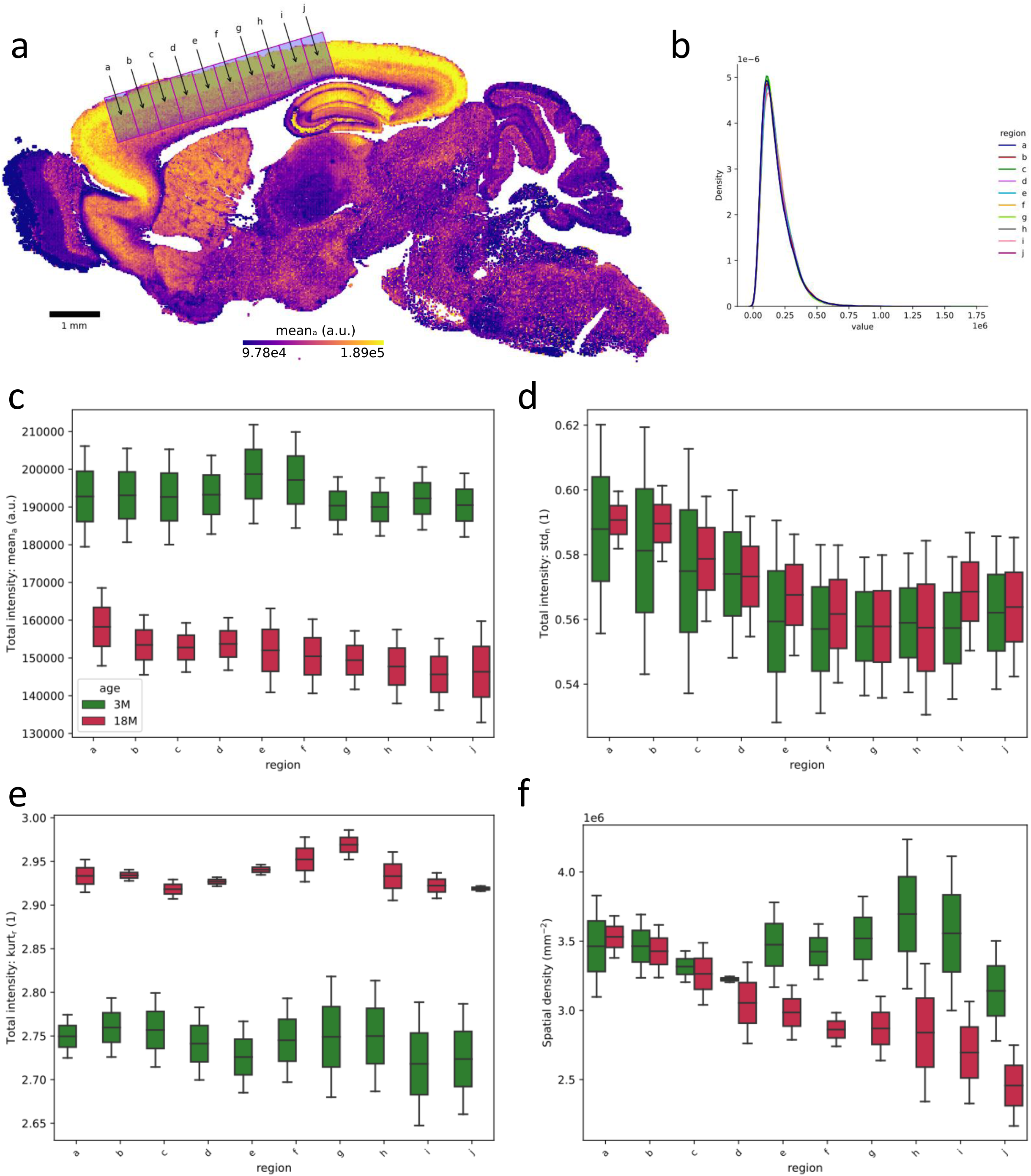
Tangential gradient (anterior to posterior) in the cortex. a. Locations of ten boxes a—j defining the gradient in a section from a three months old individual. The data in the following panels is pooled across individuals by age, where the box locations were registered across the individuals. b. Normalized total intensity histograms. c. Arithmetic means. d. Normalized standard deviations. e. Robust kurtosis. f. Spatial density; here defined as the total number of puncta per box, divided by the box size.

**Supplementary Figure S3II.**
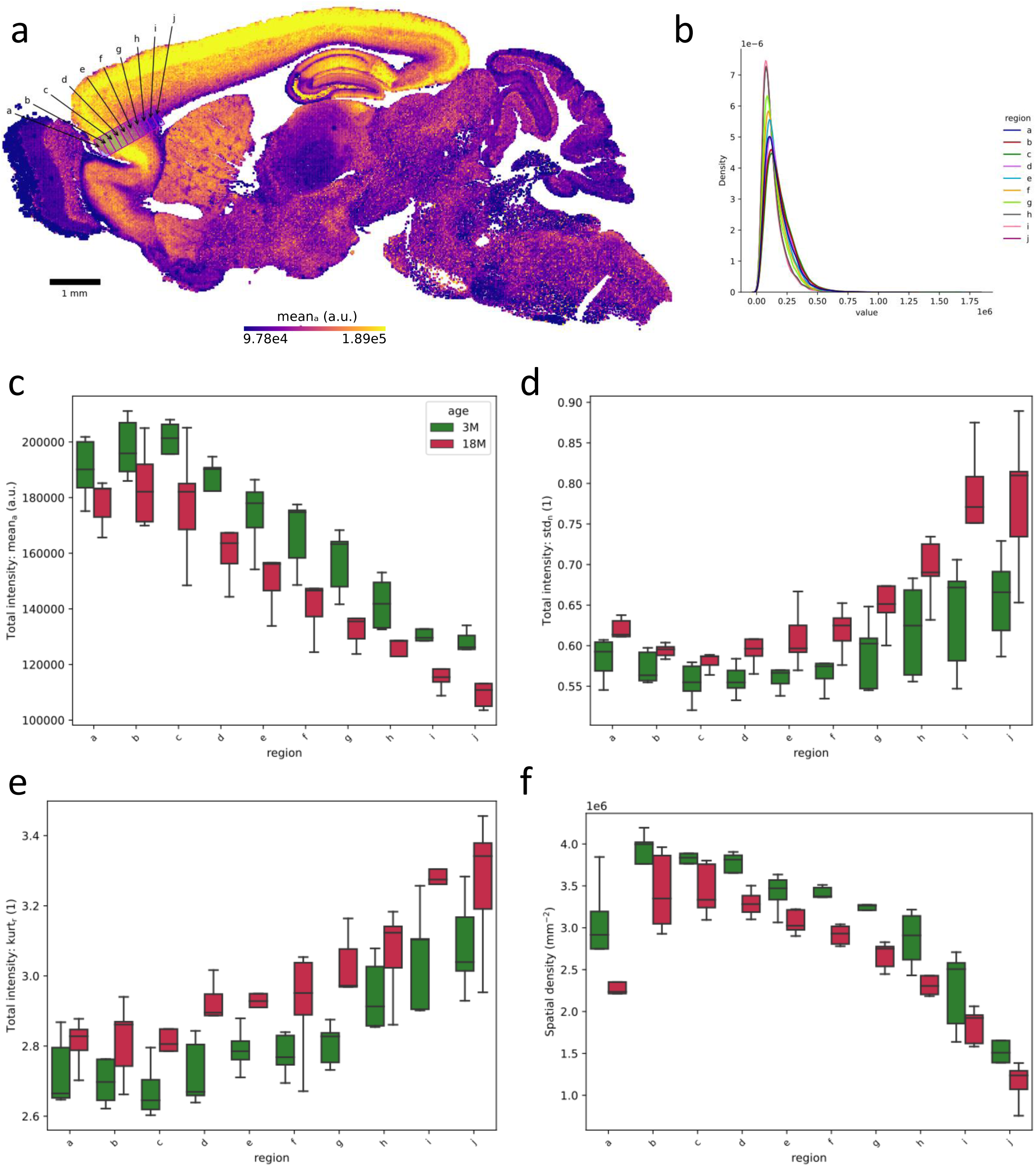
Radial gradient in the orbital cortex. a. Locations of ten boxes a—j defining the gradient in a section from a three months old individual. The data in the following panels is pooled across individuals by age, where the box locations were registered across the individuals. b. Normalized total intensity histograms. c. Arithmetic means. d. Normalized standard deviations. e. Robust kurtosis. f. Spatial density; here defined as the total number of puncta per box, divided by the box size.

**Supplementary Figure S3III.**
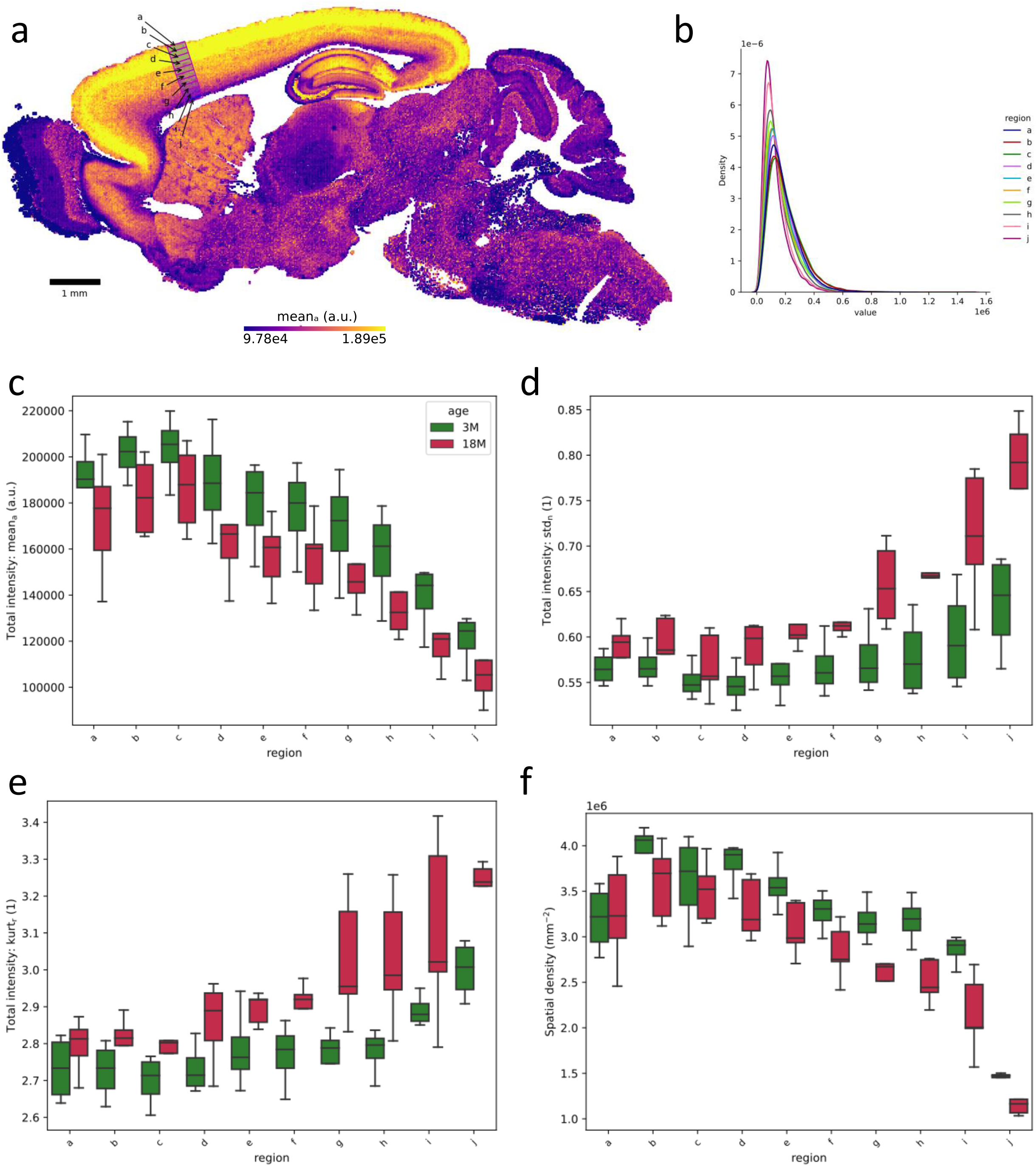
Radial gradient in the motor cortex. a. Locations of ten boxes a—j defining the gradient in a section from a three months old individual. The data in the following panels is pooled across individuals by age, where the box locations were registered across the individuals. b. Normalized total intensity histograms. c. Arithmetic means. d. Normalized standard deviations. e. Robust kurtosis. f. Spatial density; here defined as the total number of puncta per box, divided by the box size.

**Supplementary Figure S3IV.**
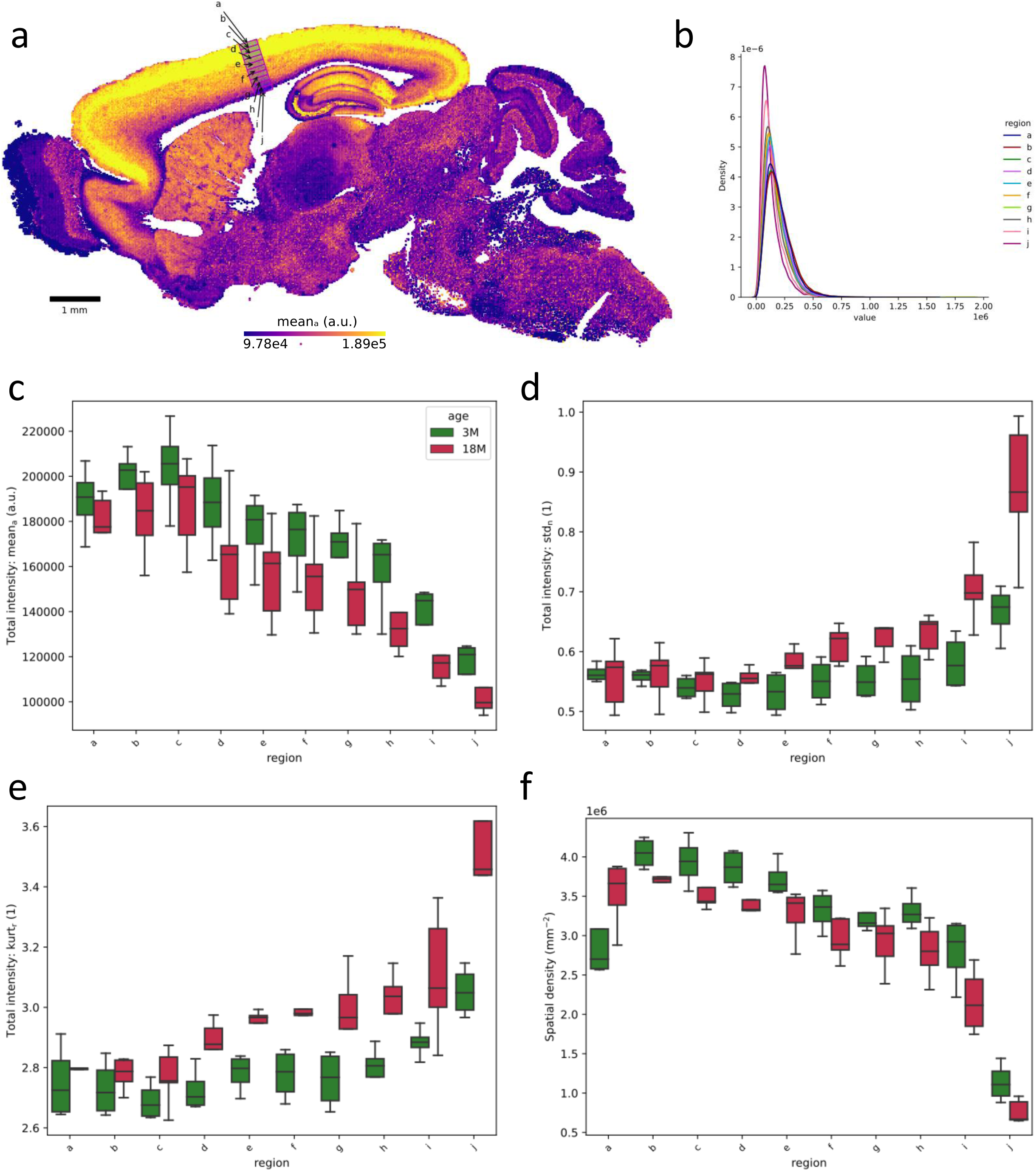
Radial gradient in the somatosensory cortex.

**Supplementary Figure S3V.**
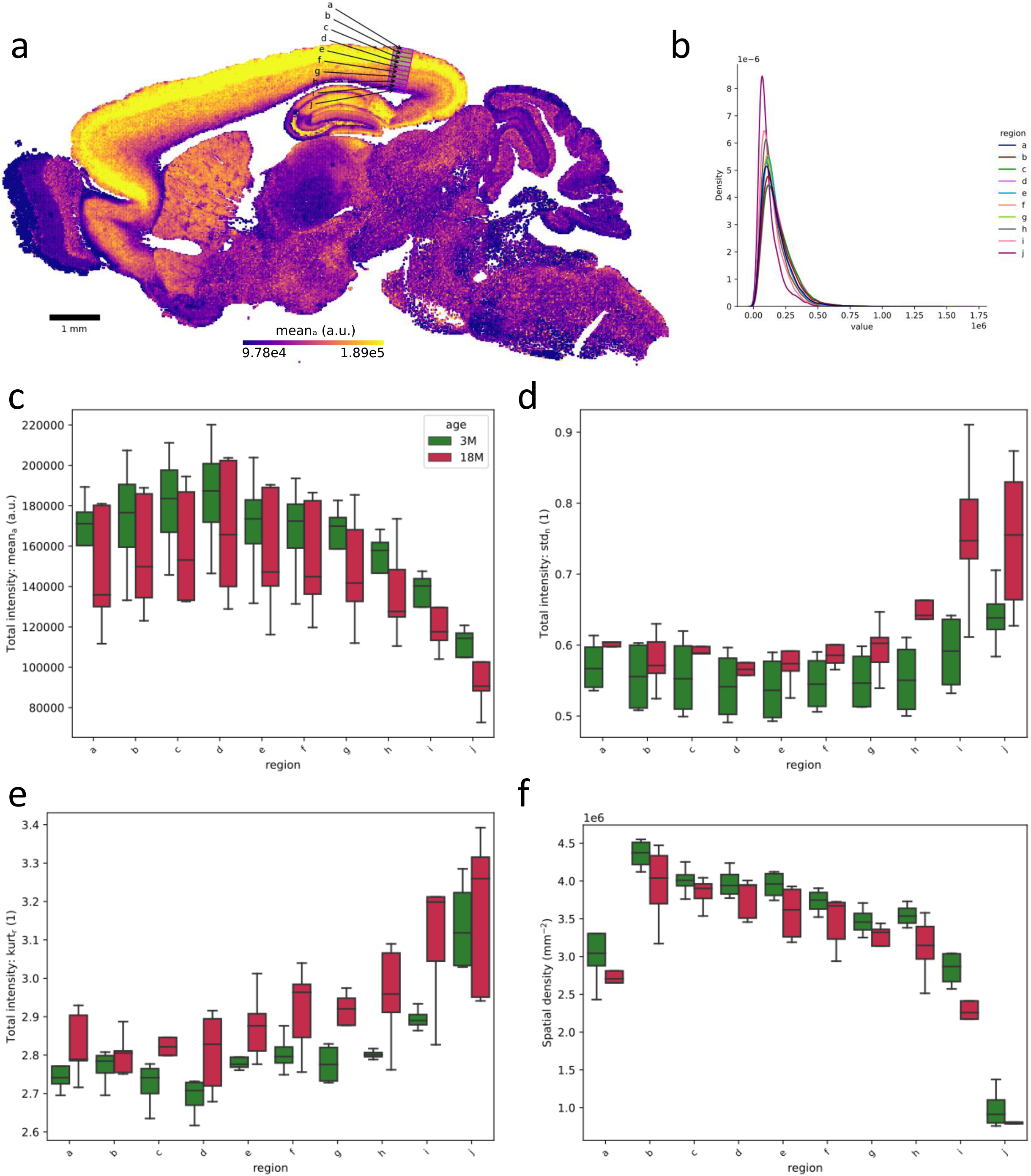
Radial gradient in the visual cortex.

**Supplementary Figure S3VI.**
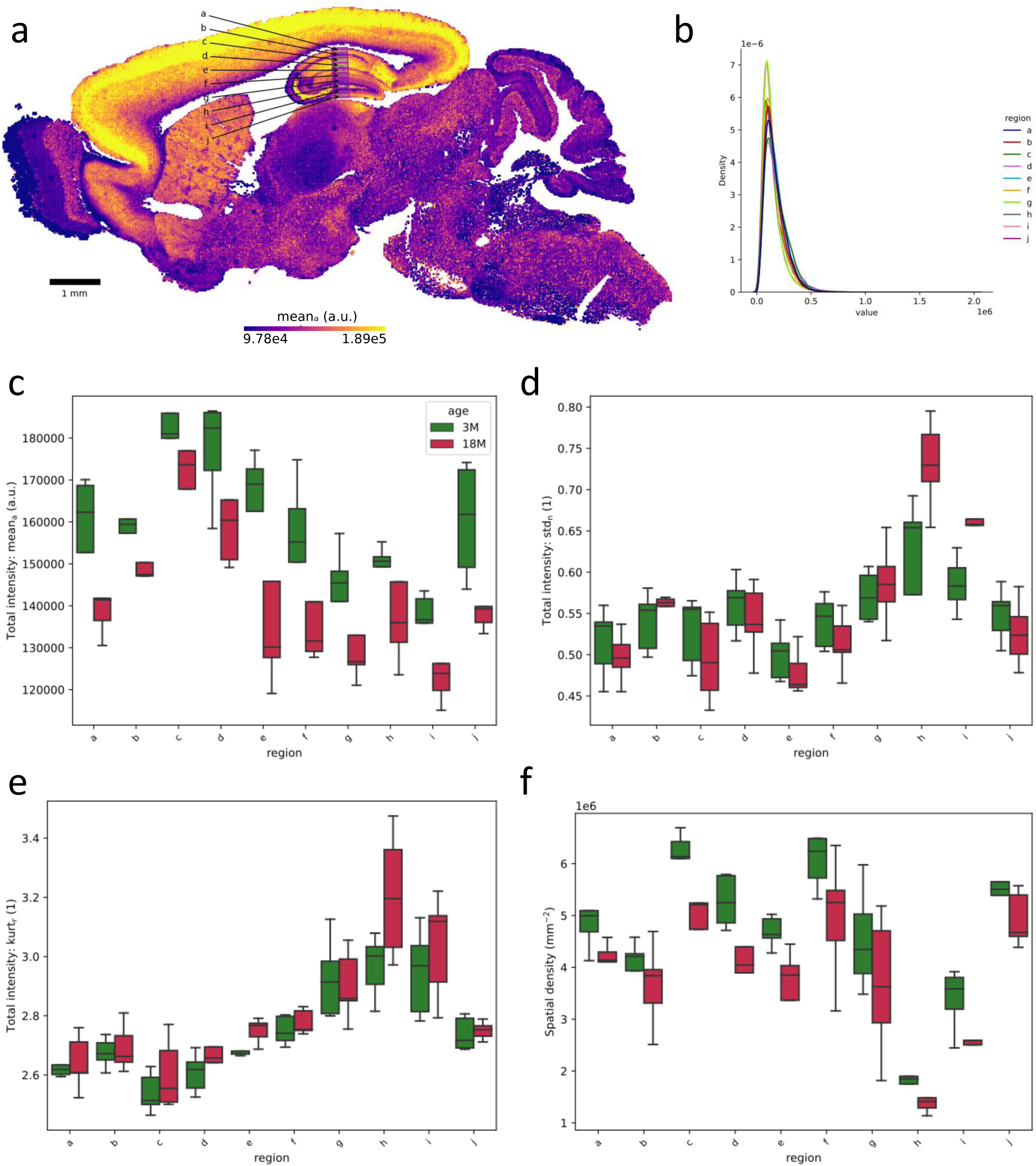
Transverse gradient (dorsal to ventral) in the hippocampus. a. Locations of ten boxes a—j defining the gradient in a section from a three months old individual. The data in the following panels is pooled across individuals by age, where the box locations were registered across the individuals. b. Normalized total intensity histograms. c. Arithmetic means. d. Normalized standard deviations. e. Robust kurtosis. f. Spatial density; here defined as the total number of puncta per box, divided by the box size.

**Supplementary Figure S3VII.**
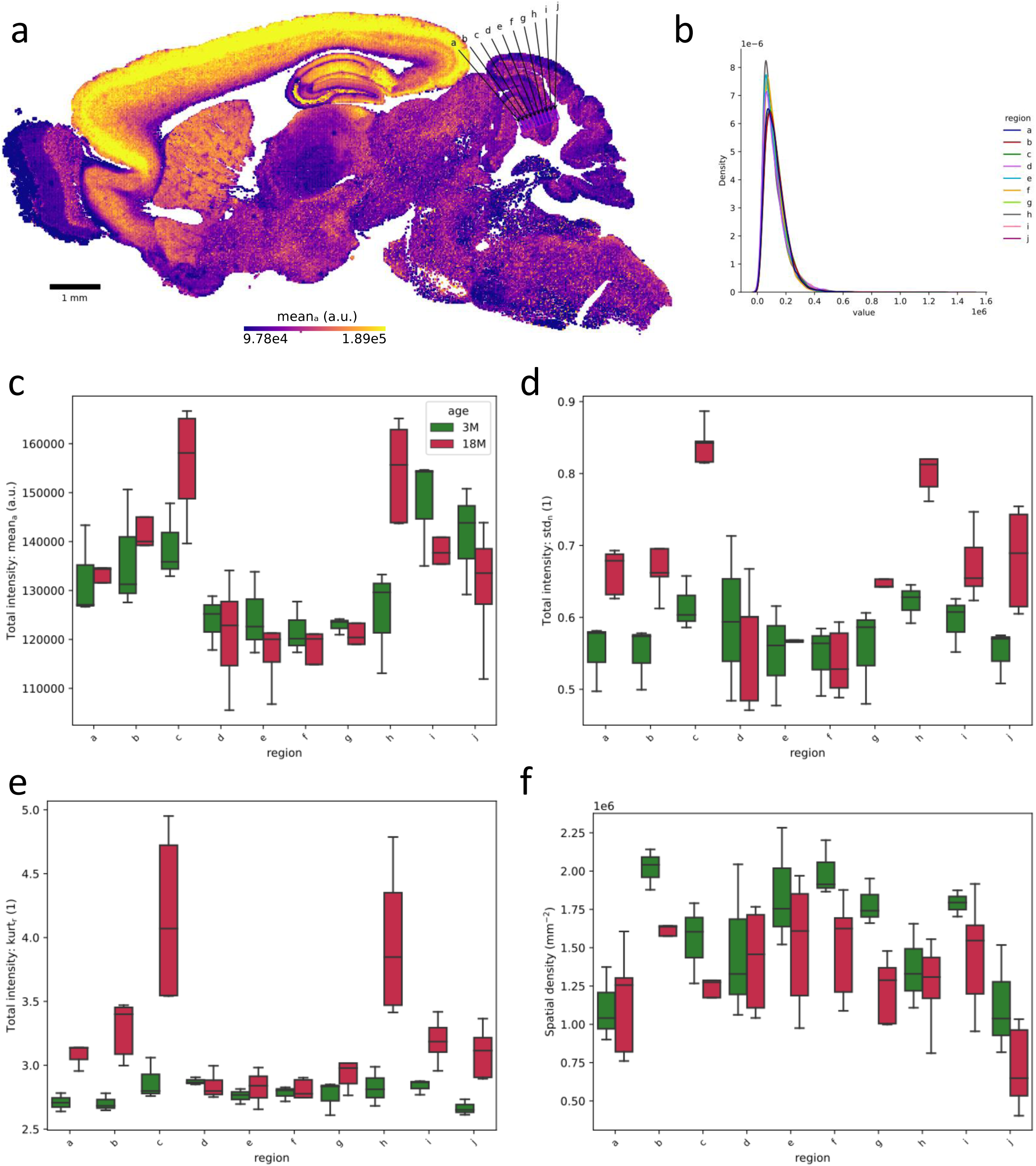
Gradient through the cerebellar layers. a. Locations of ten boxes a—j defining the gradient in a section from a three months old individual. The data in the following panels is pooled across individuals by age, where the box locations were registered across the individuals. b. Normalized total intensity histograms. c. Arithmetic means. d. Normalized standard deviations. e. Robust kurtosis. f. Spatial density; here defined as the total number of puncta per box, divided by the box size.

**Supplementary Figure S3VIII.**
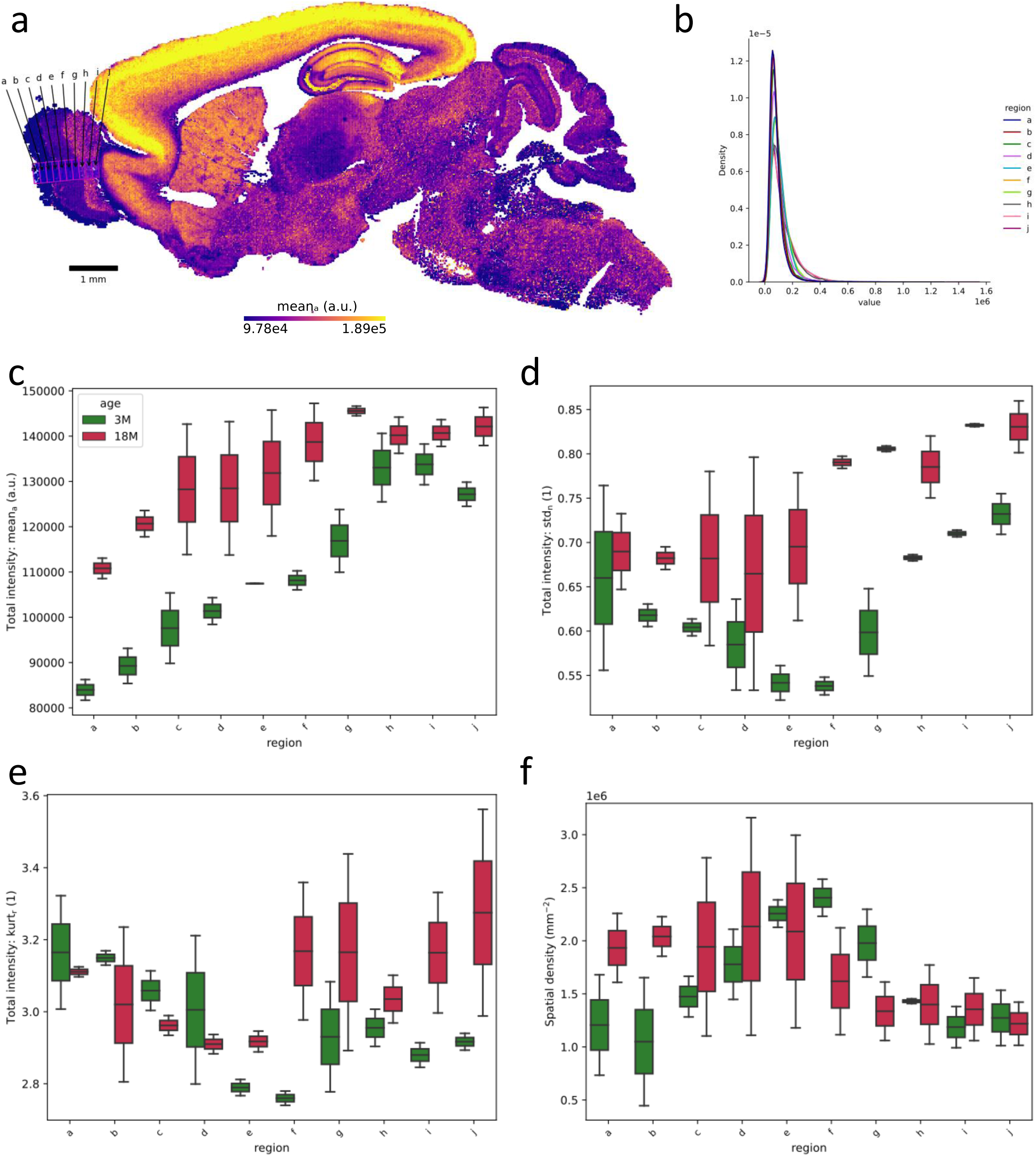
Gradient through the olfactory bulb (anterior to posterior). a. Locations of ten boxes a—j defining the gradient in a section from a three months old individual. The data in the following panels is pooled across individuals by age, where the box locations were registered across the individuals. b. Normalized total intensity histograms. c. Arithmetic means. d. Normalized standard deviations. e. Robust kurtosis. f. Spatial density; here defined as the total number of puncta per box, divided by the box size.

**Supplementary Figure S4.**
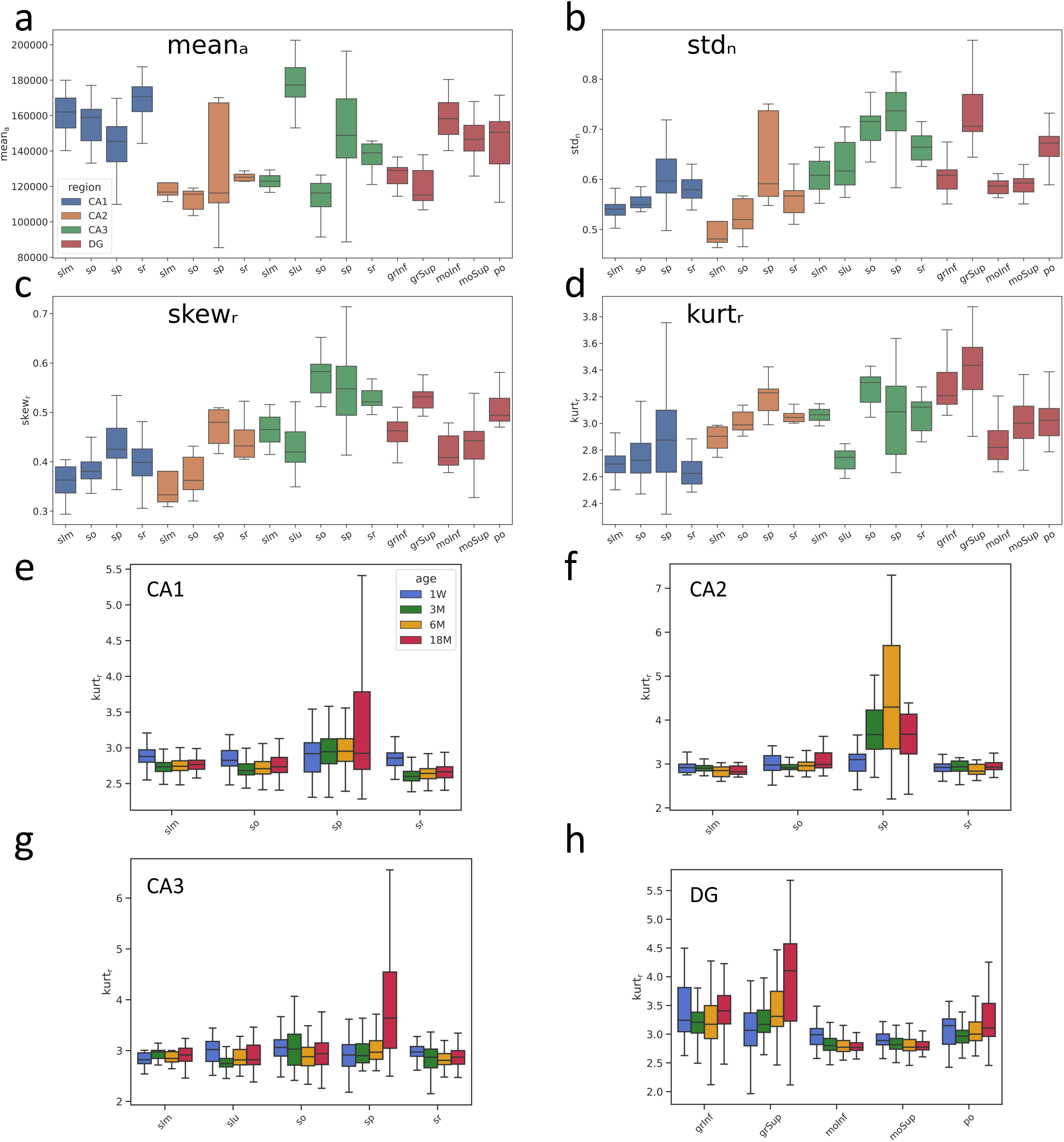
Details from the hippocampus. Bar plots of the main moments for all subregions (grouped by main constituent) of the hippocampus. a. Arithmetic mean. b. Normalized standard deviation. c. Robust skewness. d. Robust kurtosis. e. Layer-wise and age-wise robust kurtosis for CA1. f. Layer-wise and age-wise robust kurtosis for CA2. g. Layer-wise and age-wise robust kurtosis for CA3. h. Layer-wise and age-wise robust kurtosis for DG.

**Supplementary Figure S5.**
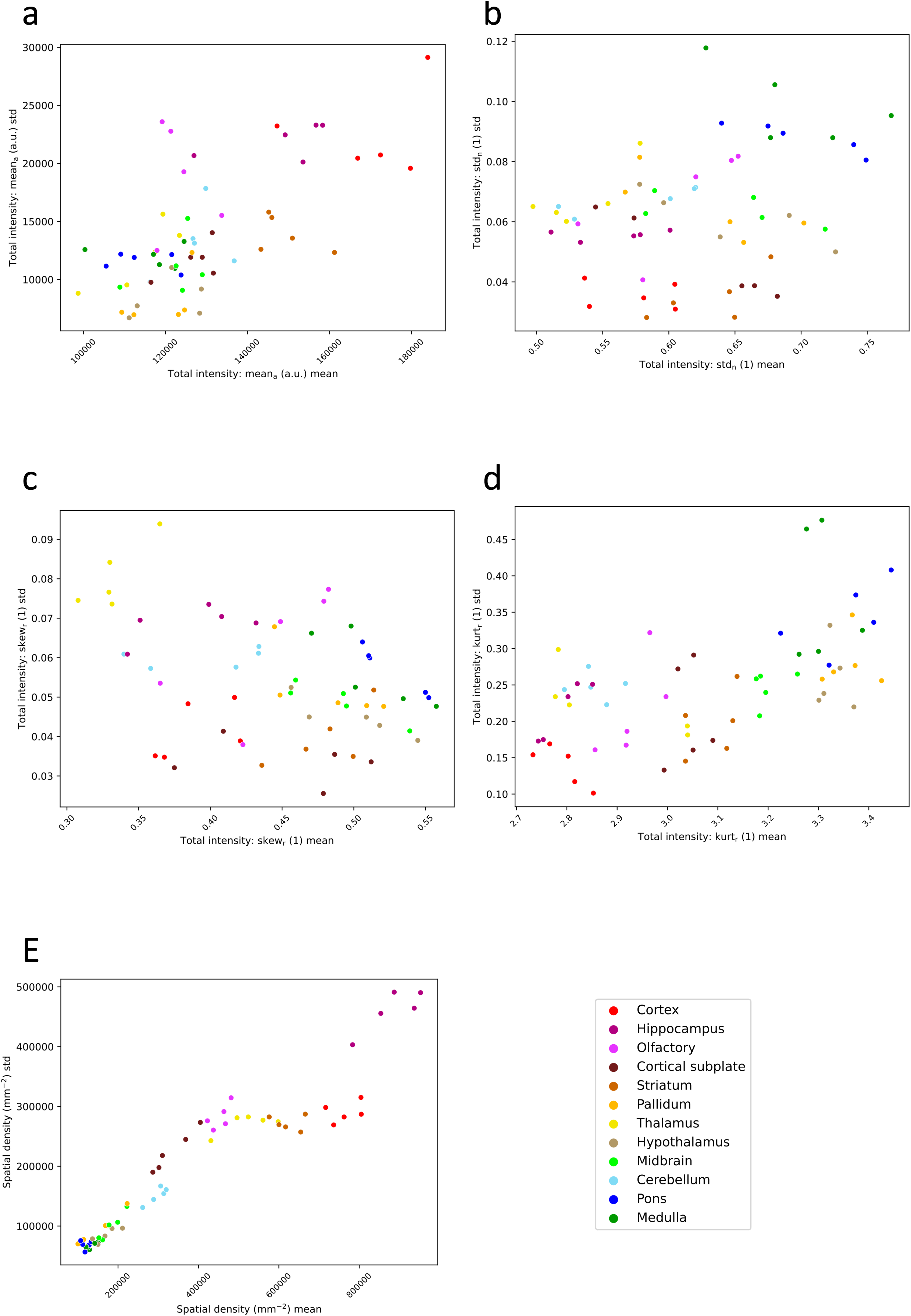
Second order moment relations. For each major region moment descriptors were computed per 100 µm x 100 µm tile, then the means and standard deviations were calculated across the tiles, and summarized per individual animal, within the three months age group. Pairwise relations between the mean and standard deviation per underlying statistical moment are shown. In each panel the mean is plotted along the x axis and the standard deviation along the y axis. Each data point corresponds to one individual. a. Arithmetic mean. b. Normalized standard deviation. c. Robust skewness. d. Robust kurtosis. e. Spatial density.

**Supplementary Figure S6.**
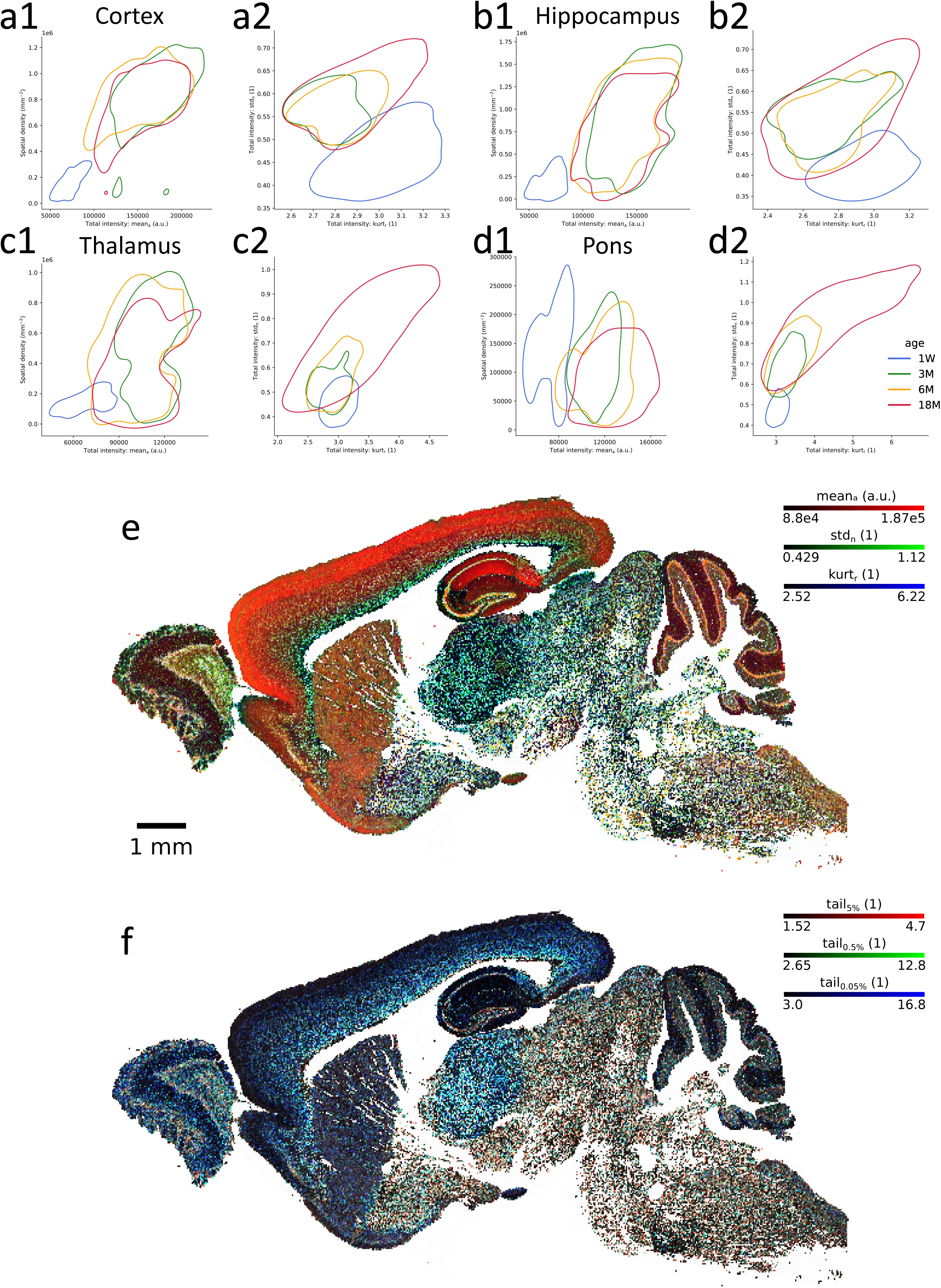
Age dependence of distribution profiles. a—d. Pairwise distributions of moments. Moment descriptors were computed per 100 µm x 100 µm tile, the values were aggregated per age group and a contour drawn around the 25th percentile. a1—d1. Artithmetic mean vs. spatial density. a2—d2. Robust kurtosis vs. normalized standard deviation. a. Cortex. b. Hippocampus. c. Thalamus. d. Pons. e—f. False color representations of triplets of moments in an 18 months old individual. Tile size 25 µm x 25 µm. Each color channel is clipped at the 5th and 95th percentiles. e. Color code by meanₐ (red), stdₙ (green) and kurtᵣ (blue). f. Color code by the relative tail values at the upper 5th (red), 0.5th (green) and 0.05th (blue) percentiles.

**Supplementary Figure S7.**
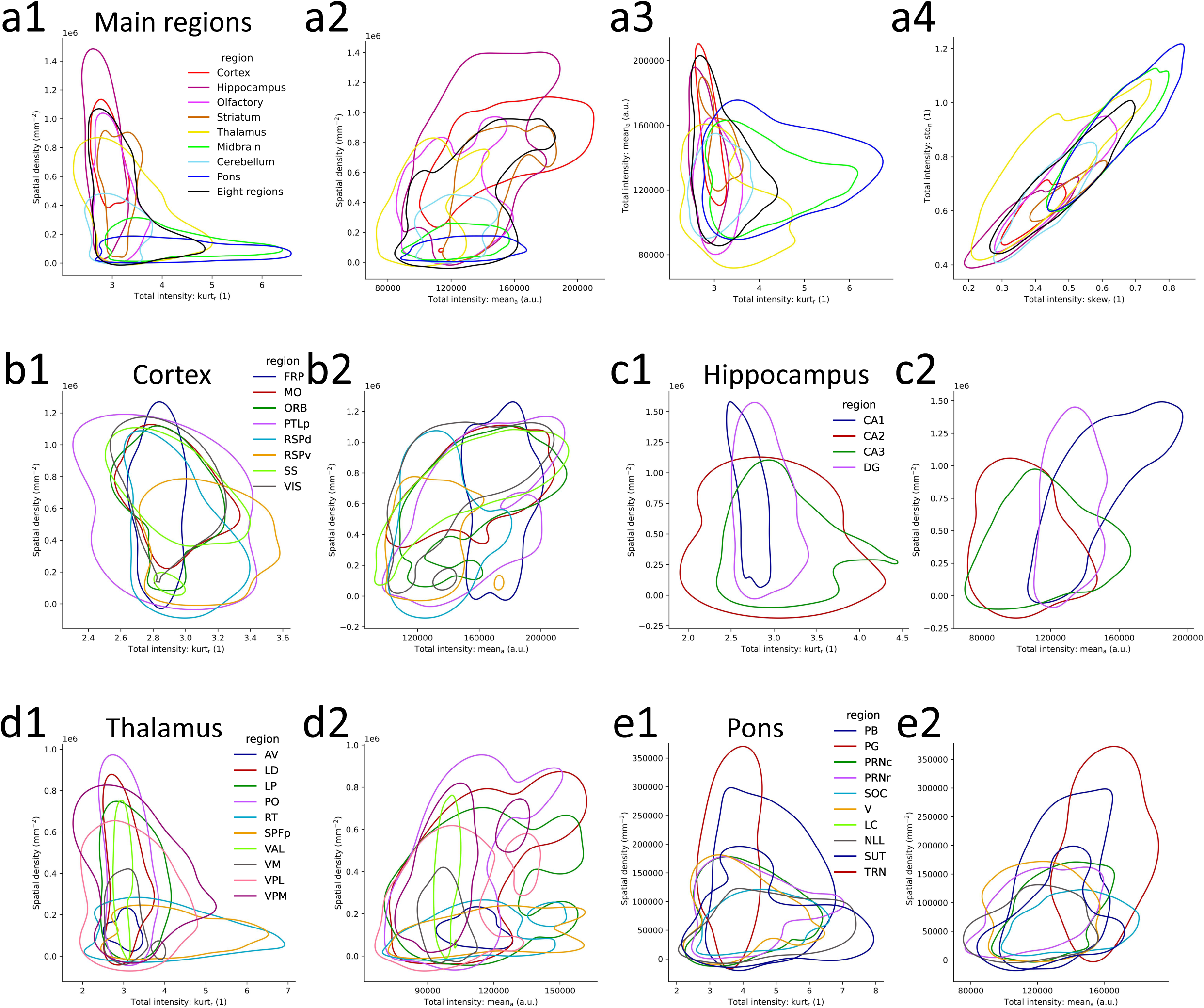
Pairwise relations of moments for the older animals. Moment descriptors were computed per 100 µm x 100 µm tile, the values were aggregated across the 18 months old animals, plotted in select pairs of moments, and a contour drawn around the 25th percentile. a1—e1. Robust kurtosis vs. spatial density. a2—e2. Arithmetic mean vs. spatial density. a3. Robust kurtosis vs. arithmetic mean. a4. Robust skewness vs. normalized standard deviation. a1—a4. Main regions. “Eight regions” refer to the pooled data across the other regions shown. b1—b2. Cortex areas. c1—c2. Main hippocampal divisions. d1—d2. Thalamic nuclei. e1—e2. Parts of the pons.

**Supplementary Figure S8.**
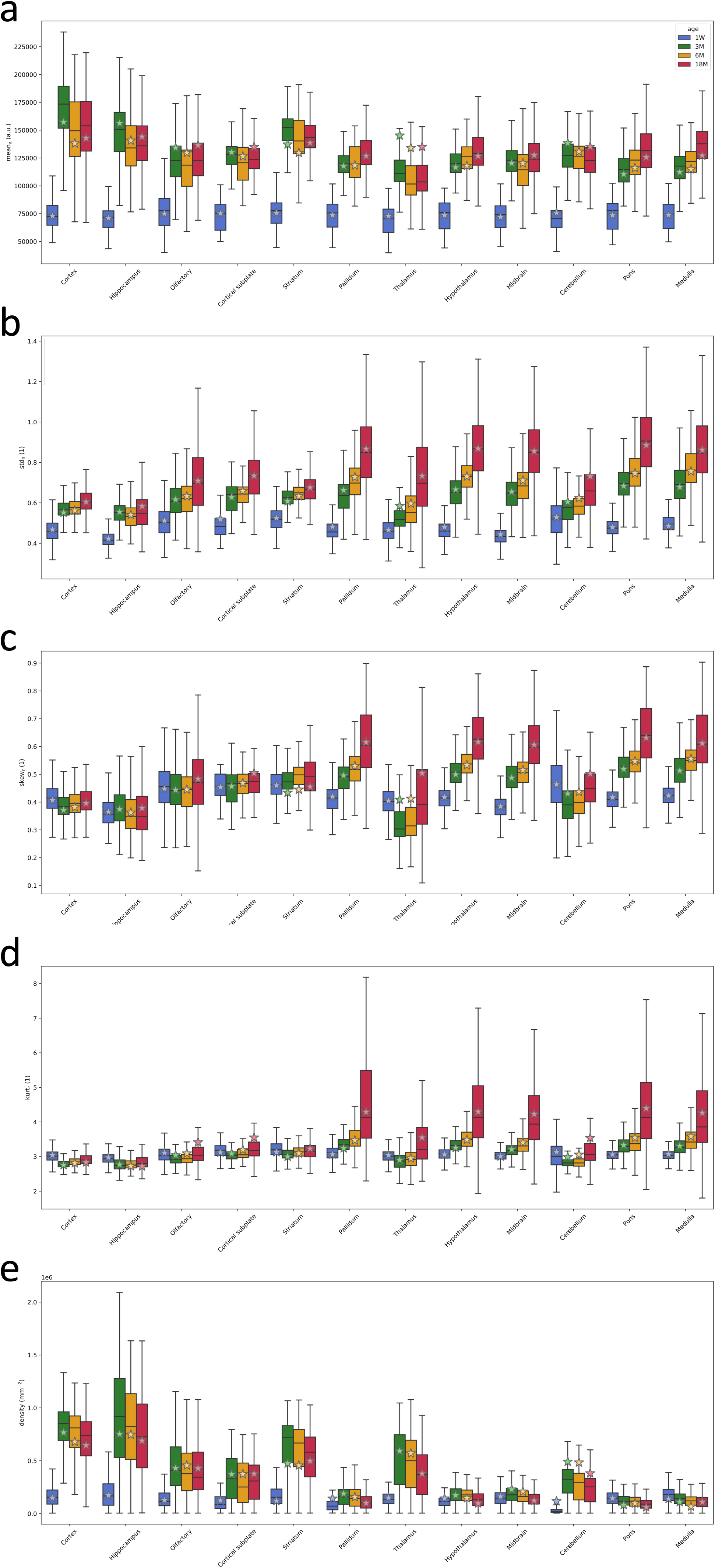
Fit of the bilinear model. The predictions of moments from the fitted bilinear explanatory model (see Methods and Fig. 6a—b). Each panel shows the predictions of a moment from the bilinear model, which was fit jointly across all five models, but independently by age. The model fit is shown by stars, whereas bars show the actual moment distributions by region, pooled from 100 µm x 100 µm tile and aggregated across individuals by age group. a. Arithmetic mean. b. Normalized standard deviation. c. Robust skewness. d. Robust kurtosis. e. Spatial density.

**Supplementary Figure S9.**
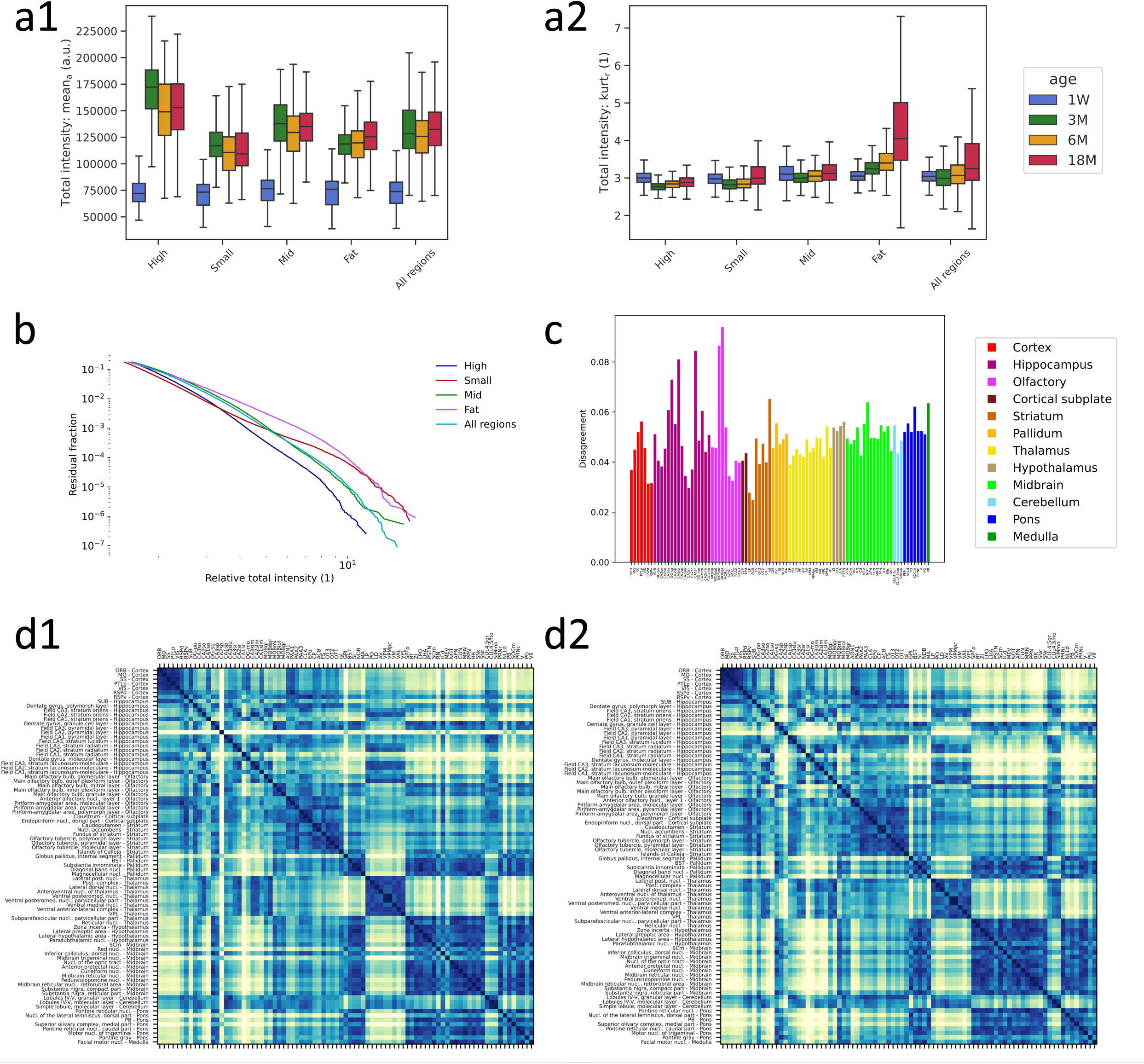
Cluster characterization (see Methods and Fig. 6e-h). a1—a2. Moment distributions by cluster. Moments were computed by 100 µm x 100 µm tile and aggregated across individuals by age group. The clustering was done for the three months age group, then mapped to the other age groups by anatomical delineation. “All regions” refer to the union of all four clusters, which is most of the brain. a1. Arithmetic mean. a2. Robust kurtosis. b. Tails of the cluster distributions. The complementary cumulative distribution function, showing the right tails (upper 15%), normalized to the respective distribution means. c. Agreement with gene expression data. Each bar displays the euclidean distance between one row (or, equivalently, column) from the distribution based distance matrix (Fig. 6e) and the gene expression based distance matrix (Fig. 6f). This is a measure of the disagreement between the two metrics when considering the estimated distance between one region and all the others. d1—d2. Distribution based distance matrices for additional age groups (compare to Fig. 6e). d1.

